# Trade-offs between risks of predation and starvation in larvae make the shelf break an optimal spawning location for Atlantic Bluefin tuna

**DOI:** 10.1101/2020.11.01.363465

**Authors:** Taylor A. Shropshire, Steven L. Morey, Eric P. Chassignet, Victoria J. Coles, Mandy Karnauskas, Estrella Malca, Raúl Laiz-Carrión, Øyvind Fiksen, Patricia Reglero, Akihiro Shiroza, José M. Quintanilla Hervas, Trika Gerard, John T. Lamkin, Michael R. Stukel

**Affiliations:** Earth, Ocean and Atmospheric Science, Florida State University, Tallahassee, FL 32306, USA; Center for Ocean-Atmospheric Prediction Studies, Florida State University, Tallahassee, FL 32306, USA; School of the Environment, Florida A&M University, Tallahassee, FL, 32307, USA; University of Maryland Center for Environmental Science, Cambridge, MD, 21613, USA; Southeast Fisheires Science Center, National Marine Fisheries Servece, Miami, FL, 33149, USA; Coorperative Institute for Marine and Atmospheric Studies, University of Miami, FL, USA; Centro Oceanográfico de Málaga, Instituto Español de Oceanografía, Fuengirola, Spain; Department of Biology, University of Bergen, Bergen, Norway; Centre Oceanográfic de les Balears, Instituto Español de Oceanografía, Palma de Mallorca, Spain

**Keywords:** Larval mortality, starvation, predation, individual based model, physical-biogoechemical model, critical period, Atlantic bluefin tuna, *Thunnus thynnus*

## Abstract

Atlantic Bluefin tuna (ABT) (*Thunnus thynnus*) travel long distances to spawn in oligotrophic regions of the Gulf of Mexico. To estimate regional larval ABT mortality, we developed a spatially-explicit, Lagrangian, individual-based model that simulates dispersal, growth, and mortality within realistic predator and prey fields during the spawning periods from 1993-2012. Modelled larval ABT experience high mortality in the first week of feeding with an average mortality rate of 0.53 ± 0.26 d^−1^ prior to postflexion. Survival ranged from 0.12%–0.32% suggesting that recruitment may vary by a factor of 2.7 due to early life stage mortality alone. Starvation is the dominant source of mortality driven by the early critical period; however, survival is ultimately limited by predation on older individuals. As a result, first-feeding larvae survive better in the more food-rich areas on the shelf, while larger larvae survive better in the open ocean with fewer predators, making the shelf break an optimal spawning area. Our findings support the hypothesis that ABT spawn in oligotrophic regions to minimize predation on their larvae. Ocean modeling tools presented in this study may help facilitate an ecosystem-based management approach to improve future stock assessment models by better resolving the stock-recruitment relationship.

## INTRODUCTION

The larval stage is a critical part of life for most fish species. A species’ pelagic larval duration is often short (days to months) relative to the life span of an individual. Nevertheless, mortality during this period can exceed integrated mortality throughout the remainder of the life cycle, with important implications for the size and health of future populations. Mortality rates typically decline with age, with initial rates >0.5 d^−1^ in some species and commonly ~0.1 d^−1^ (Houde, 2002). Survival through the egg and larval phase is often estimated to be <1% (McGurk, 1986). Consequently, larval mortality can significantly impact the number of future adults in a population and is therefore important to quantify to ensure sustainable fisheries management.

Larval mortality is a result of three main sources including predation, starvation, and losses due to advection (e.g. individuals transported away from habitat needed for settlement). Typically, predation is considered the largest source of mortality for larval fish (Peck and Hufnagl, 2012). However, depending on the species and its habitat, the magnitude of these sources can vary substantially. For example, advective losses are likely a significant source of mortality for coastal demersal species whose larvae require specific benthic substrates for settlement. In contrast, starvation is expected to be a large source of larval mortality for oceanic species like Atlantic Bluefin tuna (ABT), which spawn in warm oligotrophic regions of the Gulf of Mexico (GoM) where zooplankton biomass is low (~2–6 mg C m^−3^) and variable (Shropshire *et al.,* 2020). Because quantifying larval mortality in the field is exceedingly difficult, individual based models (IBMs) provide a strategy for investigating the relationships between larval mortality and environmental conditions. Many studies have utilized ocean models to investigate larval mortality (Hinckley *et al.*, 1996; Werner *et al.*, 1996; Heath and Gallego, 1998; Hinrichsen *et al.*, 2002). However, the majority have investigated coastal temperate species with relatively few studies of species that spawn in tropical regions.

ABT are a highly migratory species spawning offshore in the subtropical GoM from April to June (Stokesbury *et al.*, 2004). Females produce >10 million eggs (Aranda *et al.*, 2013) and individuals hatch in 1–2 days (Tanaka *et al.*, 2014). Within 2–4 days, larvae begin exogenous feeding at a size of ~3 mm length (Malca *et al.*, 2017) and ~0.1 mg dry weight (DW)(Laiz-Carrión *et al.*, 2015). The pelagic larval duration lasts 3–4 weeks (Fukuda *et al.*, 2014), during which individuals grow quickly (~0.4–0.7 mm d^−1^) (Malca *et al.*, 2017; Muhling *et al.*, 2017). Upon yolk sack absorption, larvae depend entirely on zooplankton (e.g., ciliates and copepod nauplii) ranging in size from ~100–400 μm to meet their metabolic requirements (Llopiz *et al.*, 2015; Tilley *et al.*, 2016; Shiroza *et al.,* this issue). Soon after larvae switch to feeding primarily on mesozooplankton and become increasingly piscivorous at 6-8 mm (Llopiz and Hobday, 2015; Llopiz *et al.*, 2015; Uriarte *et al.*, 2019).

The extensive literature on larval tuna provide a unique opportunity for development of IBMs. Previous studies have also identified favorable environmental conditions for ABT larvae using habitat models (Muhling *et al.*, 2010; Lindo-Atichati *et al.*, 2012; Reglero *et al.*, 2019). However, it is currently unclear if higher abundance of larvae within a spawning ground are found in certain oceanographic features because of preferential spawning, lower larval mortality rates, or physical aggregation. To investigate the impact of mortality as a mechanism for driving larval abundances in the GoM, we developed a spatially explicit, Lagrangian, individual-based model (IBM) that simulates dispersal, growth and mortality with an emphasis on the period from egg to postflexion. Larval mortalities due to starvation and predation are quantified using high-resolution, spatiotemporally-varying zooplankton fields supplied by a physical-biogeochemical model (Shropshire et al., 2020). The goals for this study were to: 1) estimate annual larval mortality; 2) compare the relative magnitudes of predation and starvation; and 3) identify environmental factors and regions in the GoM that minimize larval mortality.

## METHODS

### NEMURO-GoM model description

The IBM developed here is forced with 20 years (1993–2012) of realistic hydrodynamics, zooplankton biomasses, temperature, water clarity, and ambient light fields obtained from a three-dimensional physical-biogeochemical model (NEMURO-GoM; Shropshire *et al*., 2020). NEMURO-GoM is a highly-modified version of the NEMURO biogeochemical model (Kishi *et al.*, 2007) run in an offline configuration of the MIT general circulation model (MITgcm, Marshall *et al.*, 1997; McKinley *et al.*, 2004) and forced with daily-averaged dynamical fields from a ~4-km, data-assimilative, Hybrid Coordinate Ocean Model (Chassignet *et al.*, 2009). NEMURO-GoM was developed to examine regional zooplankton dynamics in the GoM and has been extensively validated against a combination of remote and in situ measurements including total and size-fractioned mesozooplankton biomass and grazing rates, microzooplankton grazing rates, phytoplankton growth rates, and net primary production as well as surface chlorophyll and vertical profiles of chlorophyll and nitrate. The model includes eleven state variables including two phytoplankton and three zooplankton functional groups. We briefly describe the latter, which is used to estimate the spatiotemporally-varying predator and prey fields for larval tuna. The simulated zooplankton community is composed of small zooplankton (SZ), which functionally represents heterotrophic protists (e.g. ciliates). NEMURO-GoM also includes two groups of metazoan zooplankton, including large zooplankton (LZ) that represent suspension-feeders and larger predatory mesozooplankton (PZ). These model variables are used to approximate zooplankton biomass in three size classes 0.02–0.2 mm (SZ), 0.2–1.0 mm (LZ), and 1.0–5.0 mm (PZ). For more information on NEMURO-GOM see Shropshire *et al.* (2020).

### Lagrangian model description

Lagrangian simulations were performed using the MITgcm floats package using a 4^th^ order Runge-Kutta scheme run in parallel with NEMURO-GoM. Particles (i.e. eggs) were released daily from April 1^st^ to June 30^th^ from 1993–2012 in proportion to the Bluefin Tuna Index (Domingues *et al.*, 2016), which aims to identify regions (>200 m isobath) where tuna larvae are likely to be found as a function of sea surface temperature, sea surface height, and geostrophic velocity. In total, 750,875 neutrally-buoyant, passive particles were initialized (~413 d^−1^) during the spawning period where initial depths were set randomly within the mixed layer (~5–35 m). Each particle represents a “super individual” (i.e., a group of 1000 physiologically-identical individuals that experience identical environmental forcing) (Scheffer *et al.*, 1995). The Lagrangian model interpolates three-dimensional fields of zooplankton concentration, temperature, water clarity, and ambient light estimated by NEMURO-GoM to particle positions every 6 hours. These particle attributes are used to simulate growth and mortality of larval tuna in the IBM.

The onset of piscivory in larval tuna is closely timed with the transition from flexion to postflexion stage (~2 weeks post hatch)(Llopiz and Hobday, 2015; Blanco *et al.*, 2019; Uriarte *et al.*, 2019; Laiz-Carrión *et al.*, 2019). Individuals are tracked for three weeks, fully encompassing the period when larval ABT are obligate planktivores. We focus our analysis on the period prior to when individuals switch to piscivorous behavior because NEMURO-GoM does not explicitly simulate these larger motile prey. Although the IBM does not simulate preflexion, flexion, and postflexion stages explicitly, we utilize the observed larval weight at the transition from flexion to postflexion as a reference for comparing simulated larvae to field-collected larvae. Based on larvae collected in the GoM, postflexion occurs at ~6 mm (Shiroza *et al.,* this issue) which corresponds to ~10 days post hatch (dph) and 0.54 mg DW based on relationship presented in Malca *et al.* (2017) and Laiz-Carrión *et al.* (2015). Thus, in our model simulated larvae weighing <0.54 mg DW are considered to be obligate planktivores.

### Individual-based model description

Development of the IBM in this study was guided by larval collections during two research cruises in May 2017 and 2018 as part of the Bluefin Larvae in Oligotrophic Ocean Foodwebs: Investigating Nutrients to Zooplankton in the Gulf of Mexico (BLOOFINZ-GoM) project (Gerard *et al.,* this issue). The BLOOFINZ-IBM includes three life stages (eggs, yolk-sac, and feeding larvae). Egg stage duration (h) is determined from an empirical temperature relationship (h = 4.66·exp(−0.11·θ)), where θ is temperature (Gordoa and Carreras, 2014). ABT eggs develop quickly in the warm water of the GoM and hatch in <48 hours. The probability that an egg will hatch is estimated using a temperature relationship presented in Reglero *et al.* (2018). The maximum probability of hatching (~72%) occurs at ~25°C; eggs that experience an average temperature <18°C or >33°C will not hatch. Upon hatching, particles are classified as yolk-sac larvae, and their lengths are prescribed based on a length-to-age relationship determined by Malca *et al*. (2017). In the model, length increases monotonically with age as a function of temperature (Q_10_ = 2.0), while growth in weight is dynamic (see eq1).

Although no function exists for the rate of yolk utilization for yolk sac larvae, exogenous feeding is known to begin within 2–4 dph (Tanaka *et al.*, 2014). Here, we assume exogenous feeding begins on average at 2.0 dph to be consistent with otolith-based aging studies (Malca *et al.*, 2017). The influence of temperature on the yolk-sac stage duration also uses a Q_10_ of 2.0 which results in exogenous feeding starting on average at 2.15 dph (1.5–3 dph, 95% CI). Once exogenous feeding begins, larvae are assumed to have utilized all egg yolk (i.e., there is no overlap between endogenous and exogenous feeding). Post yolk-sac larval weights are initialized based on a weight-to-age relationship determined from larvae collected in the GoM (Malca *et al.*, 2017; Laiz-Carrión *et al.*, 2015).

To simulate feeding larvae, we developed a bioenergetics model where growth in mass occurs if the assimilated fraction (α) of total ingestion (I_tot_) exceeds the metabolic requirement (R). The weight (W) of feeding larvae is updated every 6 hours using (eq1). Larvae experience starvation- and predation-induced mortality while they grow and are advected through the GoM as determined by ingestion, metabolism, starvation, and predation submodules, which are explained below.

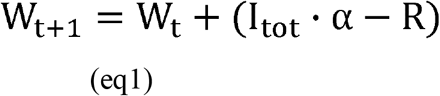

### Ingestion module

Clearance rate (m^3^ larva^−1^ d^−1^) is modeled as a function of the two-dimensional field of view fraction (φ), sensory radius (S_p_) when feeding on zooplankton prey (i.e. p = SZ, LZ, PZ), fraction of time spent feeding in a day (Δt), and the average swimming speed of larvae (v). Clearance rate is then multiplied by prey biomass (p(i,j,k,t)) at the simulated larvae’s instantaneous local position and time to estimate encounter rate (mg C d^−1^). The product of the encounter rate and capture success (σ_p_) gives ingestion rate (Ip):

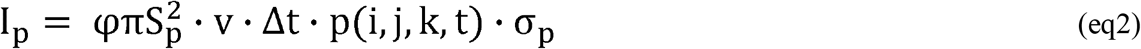

Field of view in larval fish is determined by the orientation of eyes on the head. Here, we assume that larval tuna do not perceive prey below their horizontal plane (φ = 0.5). A swimming speed of 2 body lengths s^−1^ is used for all larvae, which is approximately the average cruise swimming speed observed for cultured larval tuna (Sabate *et al.*, 2010). The number of hours larvae feed in a day is estimated as a function of latitude and day of the year. Prey biomass is derived from zooplankton biomass fields estimated by NEMURO-GoM. Unlike other pelagic larval fish, such as mackerel, which have highly-variable prey capture success through ontogeny (Hunter, 1972), the capture success for larval tuna in rearing experiments is high (>70%), even at first feeding (P. Reglero, unpub.). This is likely due to their large mouth size relative to prey (Shiroza *et al.,* this issue). Here, we assume that capture success is constant (80%). An upper limit for ingestion is set using a temperature-dependent gut turnover time with Q_10_=2.0. We use an average turnover time of 3 h at 26 °C (Young and Davis, 1990) and assume a full gut size of 10% body mass.

Ingestion is very sensitive to visual sensory radius. Hence, many mathematical formulations of sensory radius have been determined from laboratory feeding studies (Hunter, 1972), by examining the anatomy of the eye (Hilder *et al.*, 2019), or derived theoretically (Aksnes and Giske, 1993). To estimate sensory radius, we utilize a recently determined anatomical relationship for the visual acuity of larval tuna (Hilder *et al.*, 2019) along with a theoretical model of visual predation derived by Aksnes and Utne (1997) to account for the impact of light and water clarity (see online Appendix 1). This mechanistic model computes sensory radius as a function of water clarity, ambient light, larvae length, and prey size. Because larvae are primarily located in the mixed layer where light levels are high, larval length was found to be the most sensitive parameter, with sensory radius increasing monotonically with length.

### Prey field module

As larvae develop, the prey size range available to an individual changes. The shift in prey fields through ontogeny is modeled using observations of prey sizes found in the guts of larvae collected during BLOOFINZ-GoM cruises (Fig. 1A, (Shiroza *et al.,* this issue)). Specifically, we determine upper and lower bounds of prey size as a function of larval length and use this to calculate the fraction of SZ, LZ, and PZ biomass that is available to simulated larvae at their given length. We assume that PZ and LZ biomass is distributed evenly over the size range based on size-fractionated zooplankton biomass measurements (Landry and Swalethorp, this issue) and that SZ biomass follows a size spectra relationship with a slope of zero, resulting in equal distribution of biomass across equally-spaced logarithmic bins.

**Figure 1.**
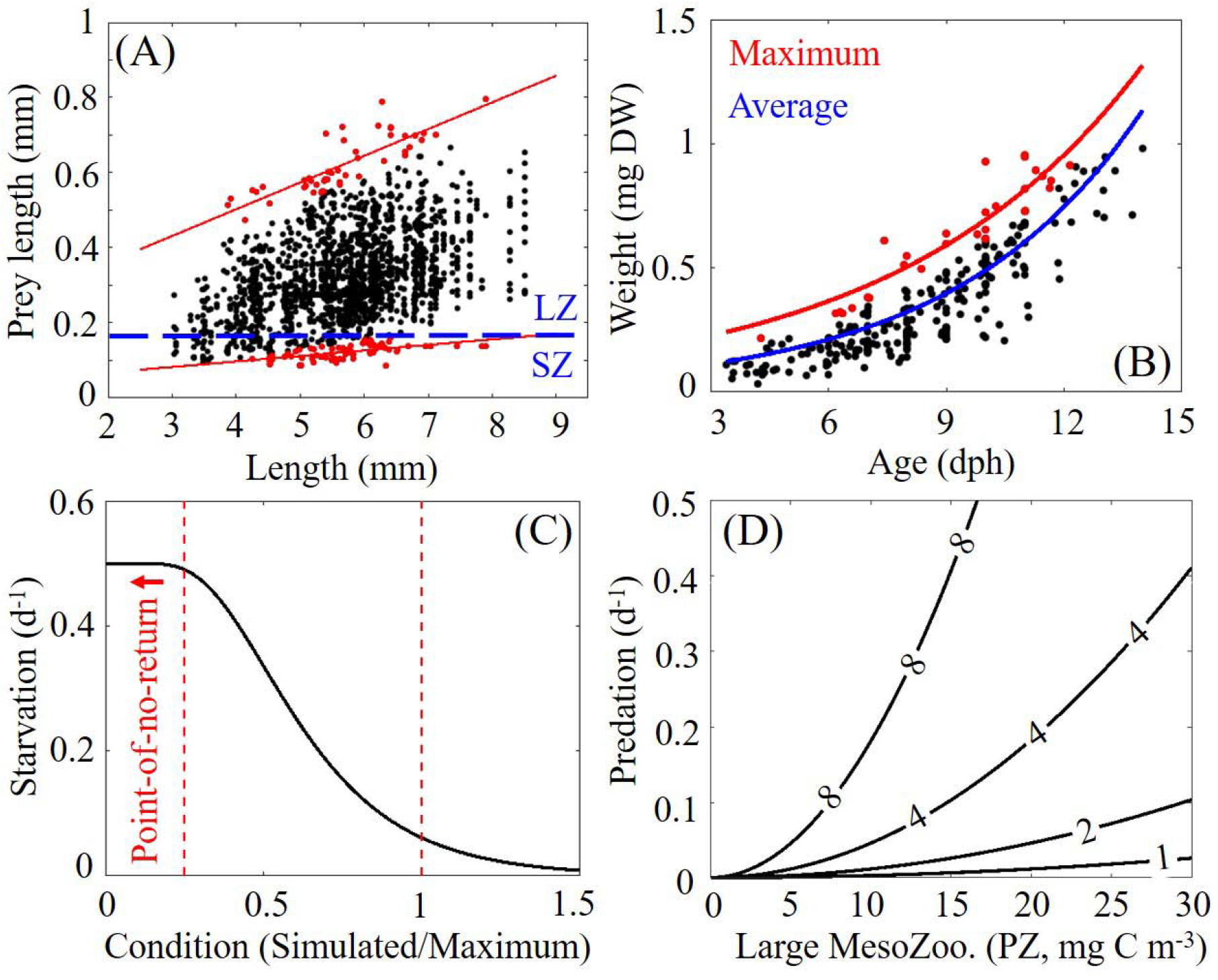
(A) Relationship between larval length (mm) and prey length (mm) from gut content analysis of 255 individuals collected in the GoM (Shiroza *et al.,* this issue). Upper and lower bounds of prey size are shown in red. Blue dotted line defines the break between zooplankton (SZ, 0.02-0.2 mm) and large zooplankton (LZ, 0.2-1 mm) NEMURO-GoM state variables. (B) Relationship between larval weight (mg DW) and age (days post hatch) for individuals collected in the GoM. (C) Starvation as a function of an individual’s condition where ≥1.0 indicates ideal condition. Condition below 0.25 is used as a threshold for the “point-of-no-return” where larvae experience irreversible starvation (increased to 1.0 d^−1^ (not shown)). (D) Predation on egg and larvae as a function of simulated large mesozooplankton (e.g. PZ, 1-5 mm) biomass and example curves of individual length at 1, 2, 4, and 8 mm.

### Metabolic requirement module

The metabolic requirement (R) of larval tuna was estimated from a weight-to-age relationship based on larvae collected in the GoM (Malca *et al.*, 2017; Laiz-Carrión *et al.*, 2015) (Fig. 1B). The derivative of this relationship gives potential growth rate in mass (dW/dA). To convert to carbon, the growth rate is multiplied by a carbon to dry weight ratio (c_f_ = 0.4; Omori, 1969). The ingestion required to meet the observed growth rate can then be estimated by dividing by an assumed gross growth efficiency (∈). Here we use a value of 0.3 (Houde, 1989). Multiplying by the difference between the absorption efficiency (α=0.7) and ∈ gives an estimate of metabolic requirement. Finally, the impact of temperature on metabolic requirement is included using Q_10_ = 2.0, yielding (eq3) where t_c_ is the temperature coefficient, θ(i,j,k,t) is water temperature at an instantaneous local position and time, and θ_avg_ represents the average water temperature that field-collected larvae experience prior to being collected (assumed to be 26°C).

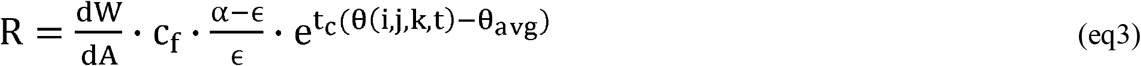

### Starvation module

To determine the probability of starvation for simulated larvae, we first identify a maximum weight at age, defined as an exponential fit to the field-collected larvae in the upper quartile of the weight-age relationship (Fig. 1B). The observed actual-weight:maximum-weight ratio is used as a metric of larval condition. We then fit a probability distribution function to the condition values for field-collected larvae and use the associated cumulative distribution function (CDF) to determine the probability of a larvae having a given condition value or lower. Finally, we perform a reflection of the CDF (i.e. 1-CDF) and scale the CDF by a maximum starvation rate parameter (0.3 d^−1^), which yields a sigmoidal function that provides a rate of mortality due to starvation given the condition (simulated-weight:maximum-weight) of a simulated larva (Fig. 1C). We prescribe elevated starvation of 1.0 d^−1^ if the simulated weight of an individual falls below 25% of the maximum-weight (i.e., condition ≤ 0.25) to account for irreversible starvation (i.e., “point-of-no-return”, (Yin and Blaxter, 1987))

### Predation module

To estimate mortality due to predation we first assume that the predators of ABT prior to postflexion (~1–6 mm) are broadly similar to the predators of the PZ zooplankton state variable in NEMURO-GoM (defined as 1–5 mm mesozooplankton). With this assumption, the mortality rate on PZ (d^−1^) estimated by NEMURO-GoM can be used to approximate predation on eggs and larvae in space and time. In NEMURO-GoM, mortality on PZ is modeled as a function of PZ biomass with a quadratic formulation to represent implicit loss to an un-modeled predator (e.g. planktivorous fish) that covaries in abundance with PZ (see online appendix 1). We assume that the abundance and predator community composition of PZ and larval tuna are identical, but that predation varies as a function of prey size, because larger prey can be detected at a greater distance. It is important to note that this further assumes that (1) escape and capture response increase proportionally as both larvae and their predators increase in size, (2) predator community is dominated by visual predators, and (3) the predator community composition does not change as larvae grow. With these assumptions (see full derivation in online appendix 1), one can compute predation-induced mortality on larval tuna (M_LT_, units = d^−1^) starting with the visual predation equation developed by Aksnes and Utne (1997) (Fig. 1D) (eq4). We note that both PZ biomass and grazing rates have been validated in NEMURO-GoM providing confidence in the PZ mortality rates that are used to approximate predation on ABT.

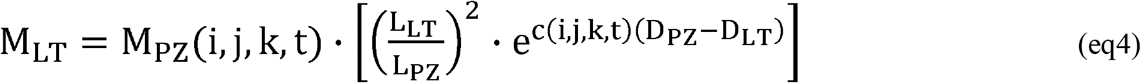

In (eq4) M_PZ_ is the specific mortality of PZ due to predation and c is the local beam attenuation coefficient (both calculated from NEMURO-GoM), L_LT_ is larvae length (during the egg stage L_LT_ is set to 1.0 mm), L_PZ_ is average length of PZ (assumed to be 2 mm), and D_PZ_ and D_LT_ are the reaction distances for predators feeding on PZ and larvae, respectively. D_PZ_ and D_LT_ are derived in online Appendix 1, although we note that the term e^c(i,j,k,t)(D^_PZ_^−D^_LT_^)^ is relatively constant and hence M_LT_ varies approximately linearly with M_PZ_ times (larval length/PZ length)^2^.

## RESULTS

### Validation of the individual based model

The BLOOFINZ-IBM was first validated by investigating larval dietary composition (Fig. 2A). In both the guts of field-collected and simulated larvae, mesozooplankton (>200 μm) constitute the majority of larval diet. Model and field measurements align with previous studies showing high dietary contributions from mesozooplankton (Young and Davis, 1990; Llopiz and Hobday, 2015; Tilley *et al.*, 2016). The dietary contribution of mesozooplankton ranged from 27% to 100% (95% CI) in field-collected larvae and 4% to 100% in simulated larvae. The majority of variability occurs in first-feeding larvae (3–4 mm size class), where mesozooplankton contributed 27% to 100% (median = 85%) for field-collected larvae and 4% to 99% (median = 59%) for simulated larvae. Dietary contribution for larvae 4–9 mm varied from 76% to 100% (median = 100%) for field-collected larvae and 66% to 100% (median = 85%) for simulated larvae. We note that while larvae are known to become increasingly piscivorous after post-flexion, only five instances of piscivory were identified in the guts of 75 postflexion larvae (5.1–8.5 mm) collected during the BLOOFINZ cruises (Shiroza *et al.,* this issue) providing further confidence in our simulated prey fields.

**Figure 2.**
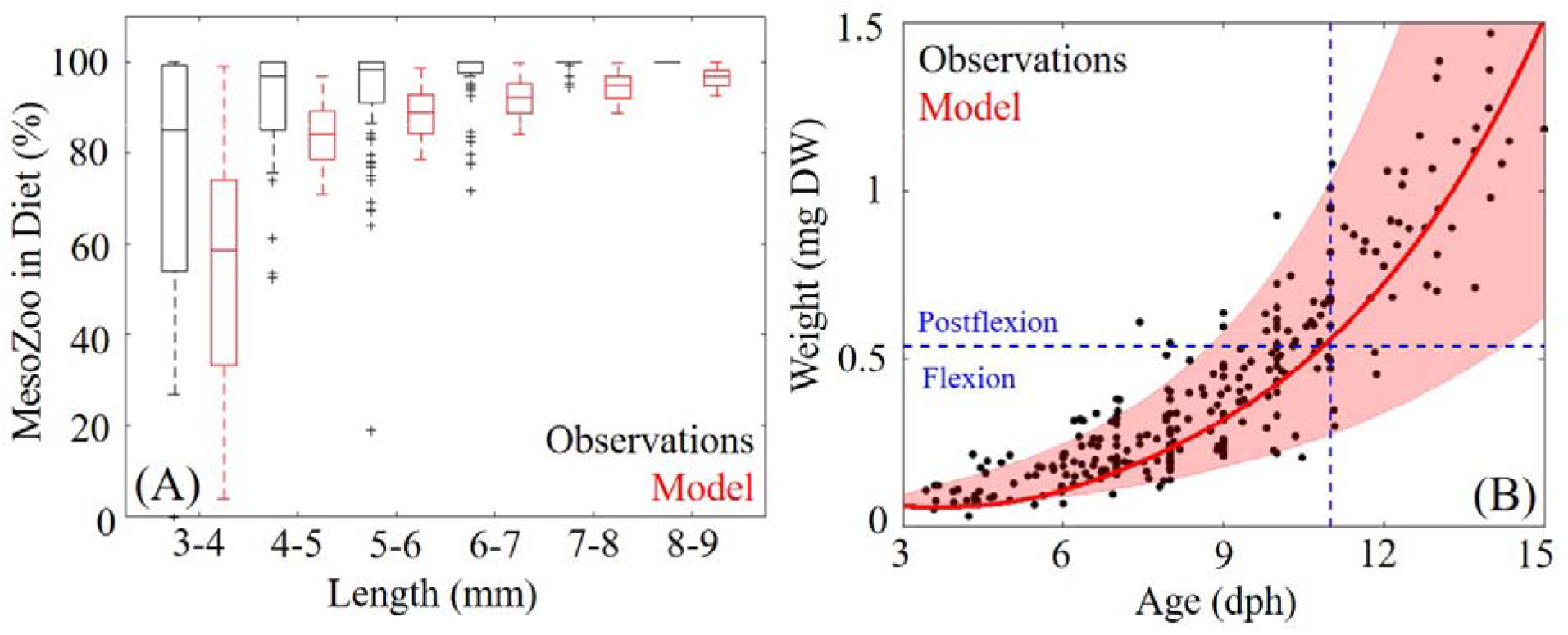
(A) Comparisons of mesozooplankton dietary contribution (% of total diet) as a function of larval length (mm) between field-collected (black) and simulated larvae (red). Whiskers extend to the 95% confidence interval. Outliers are denoted by (+) for observations and outliers for model are not shown. (B) Comparison of larval weight (mg DW) as a function of age (days post hatch) between field-collected larvae (black dots) and simulated larvae. Red line denotes model average with the 95% CI represented by shaded area. Dashed blue line denotes the average age simulated larvae reach postflexion.

Larval weights simulated by BLOOFINZ-IBM also closely match observations with a correlation of 0.94 (p<0.01) (Fig. 2B). On average, simulated larvae reached postflexion weight at 10.37 ± 1.23 dph while field-collected larvae were 10.33 dph (Malca *et al.*, 2017; Shiroza *et al.,* this issue). Herein 3 dph and 10 dph individuals are referenced as first feeding and early-postflexion larvae, respectively. The age of postflexion varied from a minimum of 8.5 dph for larvae advected on the shelf where food was abundant to a maximum of 14.5 dph (95% CI) in the most oligotrophic regions of the GoM. It is important to note that larvae may reach the average postflexion weight but have not completed the physiological development necessary to be classified as postflexion which could explain the lower bound for early postflexion age in our model. Prior to postflexion, field-collected larvae weigh 0.24 ± 0.13 mg DW while simulated larvae weigh 0.27 ± 0.13 mg DW. Although our model is expected to become more inaccurate as individuals move towards an increasingly piscivorous diet, we find nearly identical agreement between weights of simulated and field-collected postflexion larvae. On average, sampled postflexion larvae weigh 1.03 ± 0.59 mg DW while simulated postflexion larvae weigh 1.04 ± mg DW.

### Temporal variability in larval tuna survival

During the first week after spawning, the model indicates two significant mortality events (Fig. 3A). The first event involves hatching success. In our model, eggs hatch in 18–48 hours (median = 26 hours), with >28% of eggs never hatching and hence survival declines rapidly within the first two days post-spawning. Mortality slows briefly once individuals become yolk-sac larvae, with only marginally higher predation relative to eggs. On average, exogenous feeding begins at 2.12 ± 0.2 dph. Within 24 hours, the model predicts a second mortality event associated with a distinct critical period lasting ~3 days (3–6 dph). During this time, survival decreases by an order of magnitude. Survival to postflexion averaged 0.24 ± 0.05% across all 20-years of the simulation and varied from 0.12% (1995) to 0.32% (2010) (Fig. 3B). This result suggests that recruitment in the western ABT stock could vary by a factor of 2.7 due to variability in total egg and larval mortality alone. In terms of model sensitivity, we find that hatching probability, gross growth efficiency, and gut turnover/fullness parameters were the most sensitivity for determining survival to postflexion (Fig. S1).

**Figure 3.**
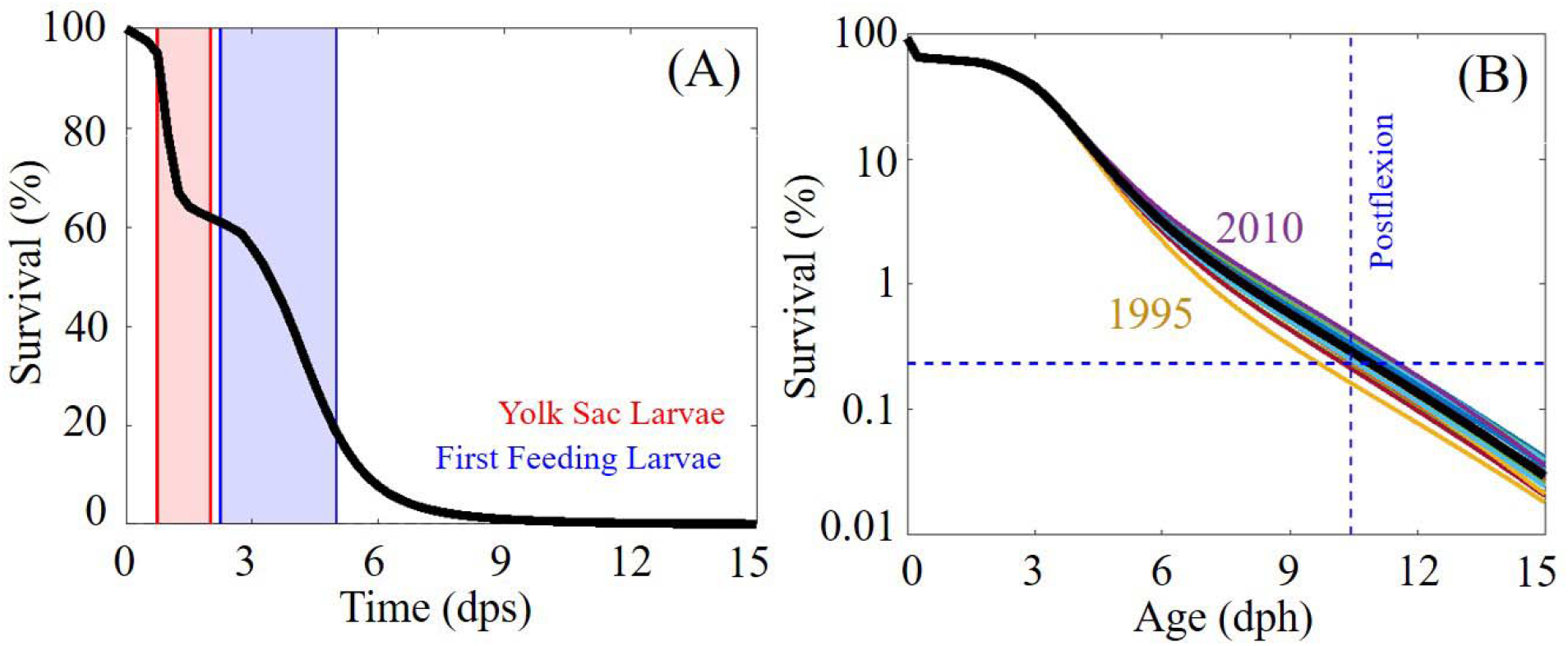
(A) Survival as a function of time (days post spawn) with red shaded area denoting yolk-sac larvae and blue shaded area denoting the period when individuals begin exogenous feeding. (B) Survival of larval tuna estimated for each year as a function of age (days post hatch) for each year (1993-2012) and black is mean of all years.

### Sources of larval mortality

Larval survival is driven by differing proportions of mortality sources as individuals develop through ontogeny. Our model indicates that starvation is the largest source of mortality prior to postflexion, accounting for 49 ± 1% of total mortality across years. Predation accounted for 20 ± 1% and mortality associated with hatching success accounted for 29 ± 1% (an additional 2 ± 0.3% was associated with advection out of the GoM). Contributions are robust even when losses are evaluated past early postflexion because of the high mortality rates during the first week. Prior to postflexion, total mortality varied from 0.06–0.93 d^−1^ (mean = 0.53 ± 0.26 d^−1^) which is slightly lower than 0.66 d^−1^ estimated by Davis *et al.* (1991). Starvation varied from 0 - 0.82 d^−1^ (mean = 0.35 d^−1^) while predation varied from 0.05–0.34 d^−1^ (mean = 0.16 d^−1^). Maximum mortality occurs at 4 dph, corresponding to the maximum rate of starvation. This result indicates that simulated larvae begin to starve <48 hours after the onset of exogenous feeding which agrees closely with results from laboratory feeding experiments of Pacific Bluefin tuna that showed ~100% mortality due to starvation within 2–4 days (Tanaka *et al.*, 2008).

To better understand why first-feeding larvae frequently starve, we investigated how prey availability evolves as larvae develop in the model. In NEMURO-GoM, SZ biomass is typically greater than LZ biomass by a factor of 3–4 in the open-ocean GoM. Hence, as larvae age and feed less on microzooplankon (SZ), they also experience a decrease in prey concentration as a result of a shift in prey size range. Prey biomass for first-feeding larvae averages 0.60 ± 0.85 mg C m^−3^ while early postflexion larvae have a prey field with 25% lower zooplankton biomass (Fig. S3F). In addition to prey fields with higher zooplankton biomass, first-feeding larvae also have lower metabolic requirements (driven by lower dW/dA). Metabolic requirement for first feeding larvae is on average 0.007 mg DW d^−1^ and increases by a factor of 7.5 for early postflexion larvae (Fig. S3A). Despite these advantages, first-feeding larvae commonly starve as estimated by our model. This is clearly a result of low clearance rates due to small sensory radii and slow swimming speeds which supports previous findings from early larval fish feeding experiments (Hunter, 1972). Clearance rates of larvae increase by more than an order of magnitude (18 L d^−1^– 480 L d^−1^, Fig. S3E) from first-feeding to early postflexion which leads to substantially lower starvation rates for larvae that survive the critical period. However, as larvae grow in our model, predation becomes an increasingly important source of mortality because their increased size allows predators to detect them more easily After this 7.75 dph, predation is the dominant source of mortality for larval ABT as estimated by our model.

### Spatial variability in starvation and predation

To quantify mortality on the shelf (<200 m isobath), the BLOOFINZ-IBM was run with random spawning (i.e. larvae were not initialized in proportion to the Domingues *et al.* (2016) larval index). Clear spatial variability in starvation and predation is predicted by the model with elevated rates of starvation in the open-ocean GoM and elevated rates of predation on the shelf (Fig. 4 B,C). Starvation is greatest in the Loop Current and the western open-ocean GoM with rates >1.5 d^−1^ and 0.5> d^−1^, respectively. The common occurrence of starvation in the Loop Current is due to warm temperatures (increased metabolic requirements) combined with low prey biomass. This result aligns with previous ichthyoplankton surveys that found low occurrences of larvae in the Loop Current (Muhling *et al.*, 2010). High starvation rates in the western open-ocean GoM can also be attributed to the Loop Current. Mesoscale eddies detach from the Loop Current every 9.5 months on average and propagate westward transporting warm oligotrophic water into the region (Sturges and Leben, 2000). Their anti-cyclonic circulation further reduces nutrient input to the surface ocean resulting in bottom-up limitation of zooplankton biomass in the western GoM (Shropshire *et al.*, 2020).

**Figure 4.**
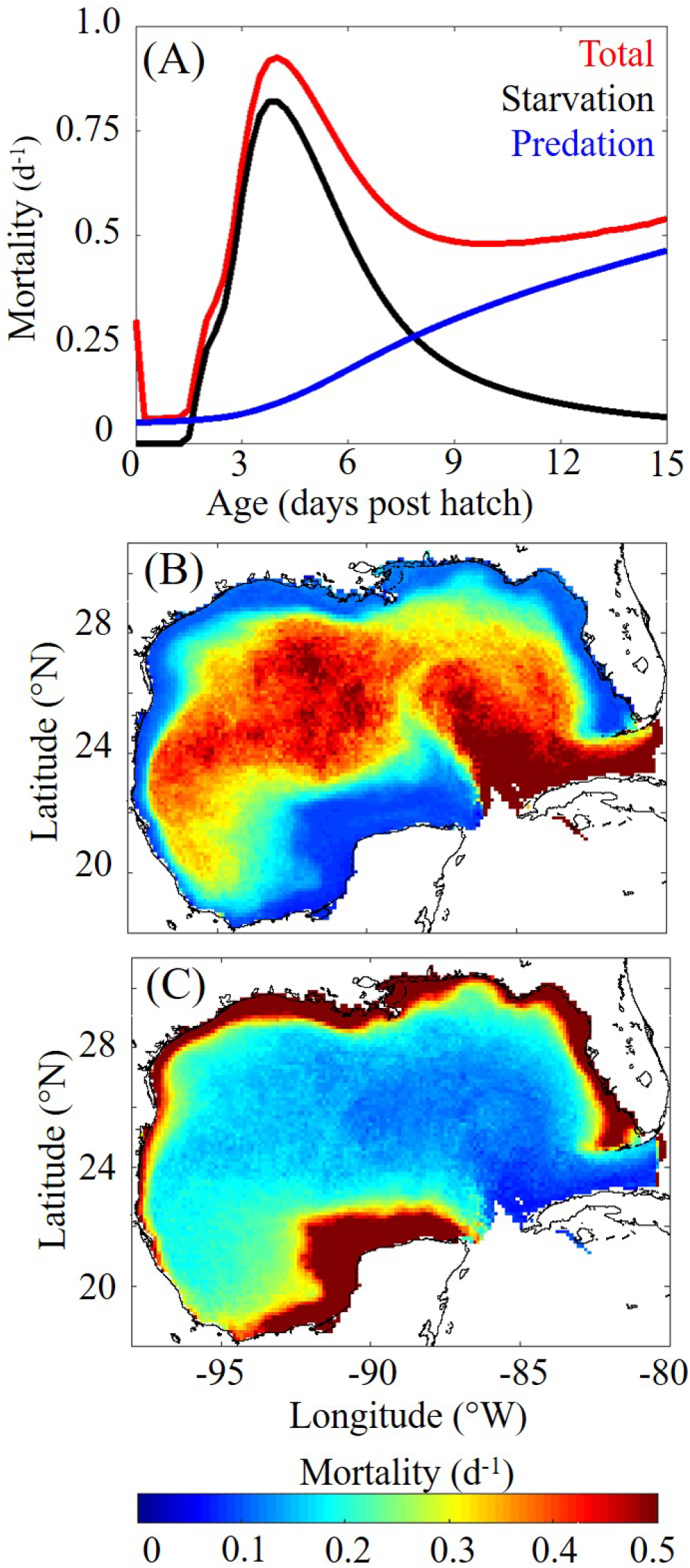
(A) Mortality rate (d^−1^) a function of age (days post hatch) with total (red), starvation (black), and predation (blue) plotted separately. (B) Spatial variability of average starvation (d^−1^) prior to postflexion. (C) Spatial variability of predation (d^−1^) prior to postflexion. Averages in starvation and predation maps are computed by organizing particles within 0.12° × 0.12° spatial bins.

Conversely, shelf regions are associated with high zooplankton biomass, allowing larvae to avoid starvation but support greater abundances of predators. Despite the decreased water clarity and ambient light on the shelf, predation increases significantly relative to the open-ocean GoM as a result of elevated abundance of PZ and hence predators. Predation rates were found to be highest near the Mississippi River and on the southern GoM shelf (Campeche Bank). In these regions, predation rates reach >6.0 d^−1^ compared to ~0.1–2.0 d^−1^ in the open-ocean GoM. High nutrient discharge near the Mississippi river mouth supports some of the highest concentrations of phytoplankton and zooplankton anywhere in the GoM (Shropshire *et al.*, 2020). This suggests that high rates of predation in this region are realistic. However, on the Campeche Bank the model may overestimate predation. As noted by Shropshire *et al.* (2020), the greatest model-data mismatch occurs on the Campeche Bank, where NEMURO-GoM commonly overestimates chlorophyll by a factor of 2–3, likely resulting in similar overestimates of PZ biomass. Hence, although predation is likely high on the Campeche Bank, our model may overestimate its magnitude.

### Spatial variability in larval survival

To investigate spatial variability in larval survival all particles were organized within 0.12° × 0.12° spatial bins based on their spawning location (Fig. 5A,B) and their time varying location (Fig. 5C,D). Both approaches reveal that the outer shelf and shelf break regions of the GoM are optimal for larval survival, minimizing the risk of starvation and predation. Larvae are particularly successful if they are spawned near the shelf break where they experience high prey concentrations during critical period and are more likely to be advected further offshore, minimizing predation as they grow. Such conditions commonly occur off the Yucatan Peninsula where upwelling stimulates phytoplankton and zooplankton biomass. Water parcels are subsequently advected offshore by the Loop Current creating a tail of plankton-rich water resulting in low starvation and predation which may explain why elevated occurrences of larvae have been documented in this region previously (Richards *et al.*, 1989).

**Figure 5.**
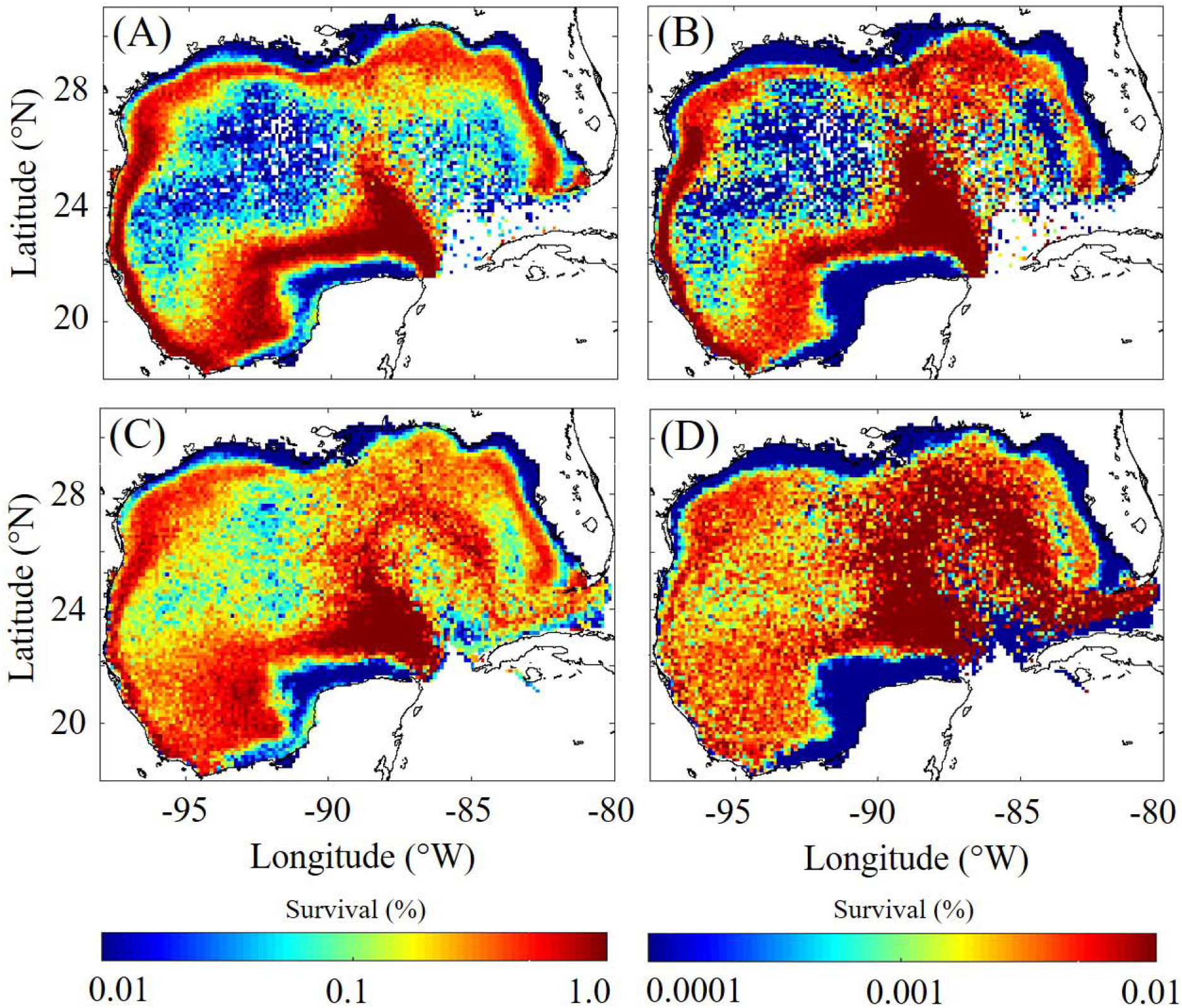
Spatial variability in larval survival to early postflexion (A,C) and late postflexion (B,D). Survival is computed by organizing particles based on their spawning location within 0.12° × 0.12° bins (A,B) and based on their time varying location (C,D).

To better understand the impact of predation on survival, which mainly influences older larvae, we investigated larval survival out to 7 days after postflexion. While our model cannot simulate prey fields of piscivorous larvae, starvation is thought to be uncommon for late postflexion larvae, as witnessed by elevated growth rates documented after the initiation of piscivory (Tanaka *et al.*, 2014). Indeed, our model estimates that starvation is substantially reduced after larvae reach postflexion. This offers some confidence that the model may provide reasonable simulations until the point where larvae develop stronger swimming behavior after metamorphosis at ~25 dph (Fukuda *et al.*, 2014). For late postflexion larvae, the model estimates that survival decreases drastically on the shelf. The highest survival again occurs at the shelf break (although shifted slightly further from shore), which suggests that while conditions on the shelf are ideal for survival of younger larvae, survival is ultimately limited by predation on older individuals. This result may explain why ABT spawn in the highly oligotrophic GoM despite the lack of food for first-feeding larvae.

The open ocean GoM is characterized by complex circulation which can significantly impact zooplankton concentrations and hence larval survival. To compare survival and mortality in different oceanographic features we organized particles into six hydrographic classifications: cyclonic and anticyclonic eddies, boundaries of cyclonic and anticyclonic eddies, common water (non-eddy regions of the open-ocean GoM), and shelf interior (<30 m isobath) (Fig. 6). To identify larvae in each of these regions, we use the dynamical criterion of Domingues *et al.* (2016) based on sea surface height and geostrophic velocity. The model estimates that survival to postflexion is lowest in the interior of anticyclonic eddies with similar survival at their boundaries (Table 1). The greatest survival occurred at the boundaries of cyclonic eddies with only marginally lower survival associated with the interiors of cyclonic eddies and common water. The greatest starvation rates occurred in the interiors of anticyclonic eddies, which were also regions with the lowest rates of predation. Starvation in the open-ocean GoM was lowest at the boundaries of cyclonic eddies and, apart from the shelf, had the highest levels of predation.

**Figure 6.**
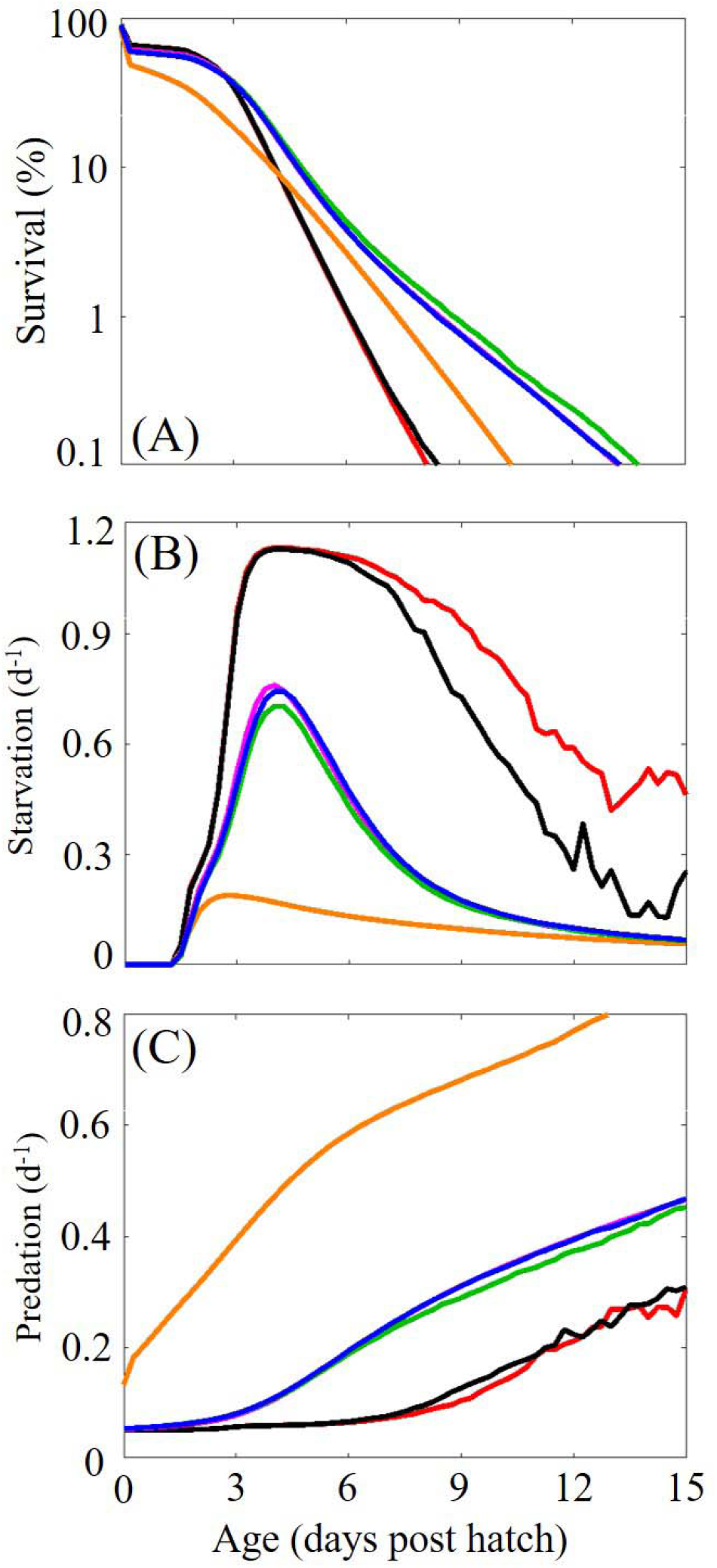
(A) Survival (%) as a function of age for larvae located inside anticyclonic regions (ACR), cyclonic regions (CR), anticyclonic boundary regions (ACBR), cyclonic boundary region (CBR), common water (CW), and shelf region (SR). (B) Starvation (d^−1^) (C) and predation (d^−1^) as functions of age for the same regions as above.

**Table 1:** Survival (%), starvation (d^−1^), and predation (d^−1^) in different oceanographic regions for early postflexion (top) and late postflexion (bottom).

|  | Region | Survival | Starvation | Predation |
| --- | --- | --- | --- | --- |
| Early Postflexion | Anticyclonic | $0.02 \pm 0.05$ | $0.74 \pm 0.14$ | $0.09 \pm 0.03$ |
| | Cyclonic | $0.98 \pm 0.27$ | $0.31 \pm 0.04$ | $0.15 \pm 0.02$ |
| | Anticyclonic Boundary | $0.05 \pm 0.07$ | $0.67 \pm 0.11$ | $0.08 \pm 0.02$ |
| | Cyclonic Boundary | $1.27 \pm 0.39$ | $0.28 \pm 0.04$ | $0.14 \pm 0.02$ |
| | Common Water | $0.99 \pm 0.16$ | $0.31 \pm 0.02$ | $0.15 \pm 0.01$ |
| | Shelf | $0.55 \pm 0.09$ | $0.10 \pm 0.01$ | $0.45 \pm 0.03$ |
|  | Region | Survival | Starvation | Predation |
| Late Postflexion | Anticyclonic | $1.7e-3 \pm 3.6e-3$ | $0.60 \pm 0.21$ | $0.17 \pm 0.06$ |
| | Cyclonic | $0.04 \pm 0.01$ | $0.20 \pm 0.03$ | $0.25 \pm 0.02$ |
| | Anticyclonic Boundary | $5.7e-3 \pm 8.1e-3$ | $0.49 \pm 0.12$ | $0.16 \pm 0.04$ |
| | Cyclonic Boundary | $0.54 \pm 0.02$ | $0.19 \pm 0.02$ | $0.24 \pm 0.02$ |
| | Common Water | $0.38 \pm 7.1e-3$ | $0.20 \pm 0.01$ | $0.26 \pm 0.01$ |
| | Shelf | $2.9\text{e-}3 \pm 1.4\text{e-}3$ | $0.07 \pm 0.01$ | $0.58 \pm 0.04$ |

### Habitat suitability

We also evaluated starvation and predation in an Eulerian framework to further characterize the GoM. Since we do not assume that past conditions influence an individual’s susceptibility to predation (i.e. the physiological condition of an individual does not impact escape response), mortality due to predation can be calculated at each grid point in the domain using the predation formulation (eq4). In contrast, starvation is a function of previous environmental forcing and hence cannot be evaluated in an Eulerian framework. Instead, to quantify susceptibility to starvation, we developed a food limitation index (FLI). The FLI is defined as the ratio of metabolic requirement to total assimilated ingestion (FLI = R / (I_tot_ ·α)), where values >1.0 indicate food limitation. These maps provide snapshots of whether a larva at a given age could satisfy its metabolic requirements at any time and location in the GoM (Fig7. A-D).

**Figure 7.**
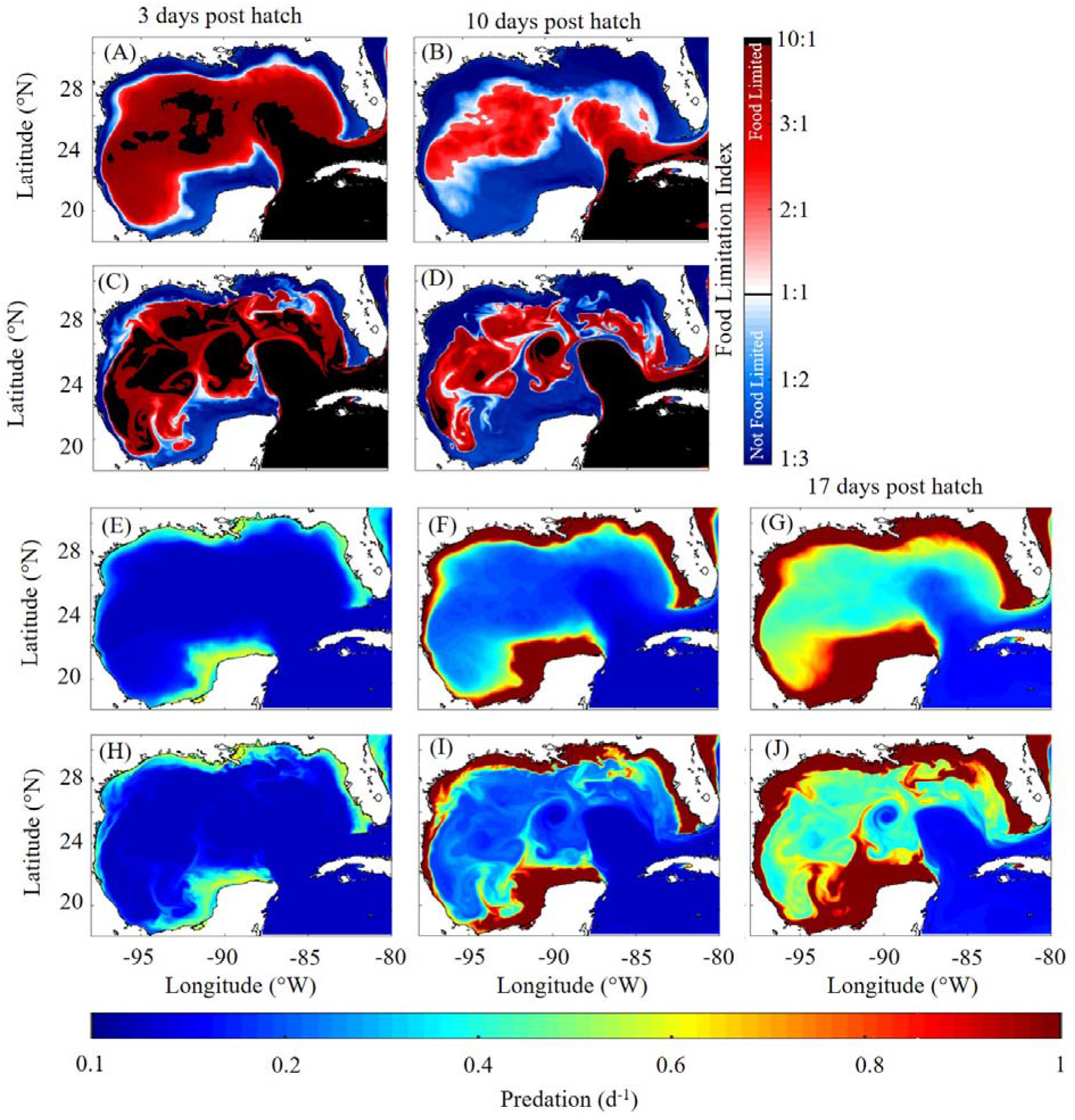
Mean and instantaneous food limitation index maps (A-D) and predation maps (E-J) for the month of May. Average food limitation index map for (A) first-feeding larvae (i.e. 3 dph) and (B) early postflexion (i.e. 10 dph). Instantaneous food limitation index map on May 15^th^ 1996 for (C) first-feeding larvae and (D) early postflexion. Average predation for (E) first-feeding larvae, (F) early postflexion larvae, and (G) late postflexion larvae (i.e. 17 dph). Instantaneous predation map on May 15^th^ 1996 for (H) first-feeding larvae, (I) early postflexion larvae, and (J) late postflexion larvae.

Daily FLI and predation maps were computed each day over the 20-year simulation during the spawning period. We find that average prey biomass in the open-ocean GoM is insufficient to meet metabolic requirements for first-feeding larvae (Fig. 7A,C). Food limitation is so severe that metabolic requirement commonly exceeds assimilated ingestion by an order of magnitude. In terms of spatial extent, food limitation for first-feeding larvae ranged from 47% to 98% of the open-ocean GoM (>200 m isobath). As evident from instantaneous fields, entrainment of shelf water by mesoscale eddies in some areas temporarily provides enough food to meet metabolic requirements for first-feeding larvae, suggesting that these events are important for the survival of larval ABT in the GoM (Fig. 7C). Consistent with starvation rates estimated by the IBM, food limitation decreases in severity and extent for older larvae. Daily food limitation for early postflexion larvae ranged from 16%–88% of the open ocean GoM and is typically confined to the Loop Current and GoM interior where Loop Current eddies are common (Fig. 7B).

The spatial extent of food limitation increased from April to June for both first-feeding and early-postflexion larvae driven by decreased prey biomass. Increased temperature later in the spawning season had an approximately neutral impact on food limitation because larvae grew (in length) faster which increased their clearance rates but also had greater metabolic requirements. On average 87 ± 7% (48 ± 15%) of the open-ocean GoM in April, 92 ± 2% (69 ± 8%) in May, and 95 ± 2% (79 ± 4%) in June is food limited for first-feeding larvae (early postflexion larvae). Predation maps show the expected inverse relationship, with elevated predation on the shelf relative to open-ocean regions (Fig. 7 E-J). On the shelf, predation varies from 0.05–0.60 d^−1^ (median=0.23) for first-feeding larvae (Fig. 7E,H), 0.14 to 1.95 (median=0.76) for early postflexion larvae (Fig. 7F,I), and 0.33 to 4.45 (median=1.76) for late postflexion larvae (Fig. 7G,H).

## DISCUSSION

ABT are highly selective spawners with adults traveling long distances from feeding grounds in the North Atlantic to spawning grounds in the GoM (Block *et al.*, 2001). Within the GoM, adult ABT are thought to spawn preferentially in specific oceanographic features (Lindo-Atichati *et al.*, 2012; Wilson *et al.*, 2005). In contrast to tropical tunas, ABT spawn over short periods of 6–8 weeks (Muhling *et al.*, 2010). Hence, the health of the western ABT stock is likely sensitive to annual environmental conditions in the GoM. Understanding how environmental conditions impact larval survival may help to better resolve the stock-recruitment relationship and is an important step for ecosystem-based management of ABT. Mesoscale variability and the resulting spatiotemporal extent of favorable larval habitat has been estimated for larval ABT by previous studies (Domingues *et al.*, 2016; Muhling *et al.*, 2013; Reglero *et al.*, 2014). However, the underlying mechanisms that make specific oceanographic features more favorable than average ambient waters have not yet been identified. Investigating the impact of larval mortality as a mechanism for driving larval abundances in the GoM was the primary objective of this study.

### Model-data misfits

The BLOOFINZ-IBM successfully resolves key dynamics pertaining to larval ecology of ABT, including realistic larval diet and weight as a function of age, stage duration, required time for the onset of starvation, and a distinct critical period associated with elevated mortality that aligns with theory (Hjort, 1914). However some model-data discrepancies exist. The model slightly overestimates the contribution of microzooplankton to larval diet across all size classes. This discrepancy may result from poor preservation of soft-bodied microzooplankton (e.g., aloricate ciliates) in fish gut contents, leading to an underestimate in the field data. Alternately, this model-data mismatch may arise from an overestimation of SZ biomass by NEMURO-GoM or errors in the IBM ingestion formulation. Simulated larvae have strict size-thresholds for prey availability that change with age, but are otherwise not selective. However, optimal foraging theory suggests that when multiple prey types are available, larvae should preferentially feed on larger, more calorie-rich prey items (Crowder, 1985; Barnes *et al.*, 2010). Indeed, Shiroza *et al.* (this issue) found that larvae were more selective for appendicularians and podonid cladocerans when these taxa were more abundant. Further realism could be added to our ingestion formulation by incorporating optimal foraging decisions (Visser and Fiksen, 2013).

Model estimates of larval weight were found to agree closely with observations, even after early postflexion, when larvae are known to become increasingly piscivorous. However, during the first few days of exogenous feeding (i.e. 3–6 dph), the model notably underestimates larval weights. On average, simulated larvae were 31% lighter relative to specimens collected in the field (data: 0.13 ± 0.05 mg DW vs model 0.09 ± 0.01 mg DW). This discrepancy may be due to the fact that in the model endogenous and exogenous feeding does not overlap although, in reality, larvae may feed exogenously while still utilizing their yolk sac. Furthermore, processes such as micro-scale turbulence or prey motility that can increase encounter rates under some circumstances are not included in our model (MacKenzie *et al.*, 1994)(Fiksen and MacKenzie, 2002). Such process may be particularly important for weakly-swimming first-feeding larvae and could be included in future versions of BLOOFINZ-IBM.

### Regional and mesoscale spatial variability in larval survival

The region of maximum survival to postflexion and late postflexion estimated by the model occurs consistently near the shelf break. This result broadly agrees with patterns of larval occurrence determined by net tows (see Fig. 3 of Muhling *et al.*, 2017), although the region of maximum survival estimated by our model is slightly closer to shore. This difference suggests that predation on the shelf may be higher than estimated here. Alternatively, ingestion may be too high on the shelf because explicit prey-dependent handling time is not included in our model or because light attenuation is underestimated leading the model to underestimate starvation.

Tagged adult ABT prefer lower continental slope waters (Teo *et al.,* 2007a) which closely aligns with the region of maximum survival predicted by our model. The presence of adults near the shelf break could indicate that elevated spawning occurs in this region. Preference by adult ABT for lower continental slope water has also been documented in spawning grounds in the Mediterranean Sea (Garcia *et al*., 2005; Alemany *et al*., 2010), and in other large pelagic species such as adult swordfish (Tserpes *et al*., 2008). This suggests that the region of maximum survival predicted by our model may be robust across regions and applicable to other large pelagic species with spawning grounds that overlap with ABT.

Our model also aligns with previous studies predicting the lowest survival inside anticyclonic eddies (Muhling *et al.*, 2010; Lindo-Atichati *et al.*, 2012) although, in contrast to previous results, it suggests the boundaries of anticyclonic eddies are only marginally better habitat. Thus, based on our model, we conclude that high occurrence of larvae in these regions is not due to lower mortality and, if borne out by future field data, must be driven by aggregation due to convergent flow or preferential spawning. However, we caution that our model may underestimate zooplankton biomass at the boundaries of anticyclonic eddies if submesoscale dynamics, which are not fully resolved by hydrodynamic models, are an important mechanism for supporting elevated zooplankton biomass.

As demonstrated by our model, patches of high prey abundance in the open ocean can be ideal spawning habitat for ABT, because these conditions can minimize starvation for first-feeding larvae, while avoiding the high predation that occurs on the shelf. Such conditions commonly occur due to entrainment of shelf water by mesoscale features. The large shelf region in the GoM and an active eddy field that drives cross shelf exchange could be an important regional characteristic that explains why ABT spawn in the GoM and not in other nearby regions with similar temperature and zooplankton biomass.

### Mortality sources through ontogeny

Mortality during the pelagic larval duration was highly variable with a distinct critical period associated with a peak in mortality driven by elevated starvation at ~4 dph. This period was followed by an ~8-day decline in mortality and then a slow rise in mortality attributable to increased predation. Increasing predatory risk could potentially be extrapolated until larvae reach metamorphosis at ~22 mm. Once larvae transition into the juvenile stage behavior substantially changes, including behaviors that may reduce predation. Relatively little is known about juveniles until they are 1–5 years old and caught in and around the Gulf Stream (Galuardi and Lutcavage, 2012). However, in laboratory experiments, late postflexion larvae and juvenile Pacific bluefin display schooling behavior as early as 25 dph (Fukuda *et al.*, 2014; Sabate *et al.*, 2010). Hence, early onset of schooling suggests that predation is likely a significant source of mortality for older larvae, as suggested by our model.

### Application to stock assessments and future work

Stock assessment models are based on population dynamics and driven by demographic data of mature fish populations. Historically, environmental forcing has largely been neglected in these models. While the mature portion of a stock is strongly driven by population dynamics, individuals in the larval stage are also significantly tied to environmental dynamics because they are planktonic. Therefore, classic age-structured modeling approaches in isolation are not well suited to model age-0 individuals. Indeed, stock-recruitment relationships (SRR) often break down, with these model-data mismatches attributed to factors outside of spawning stock biomass (Subbey *et al.*, 2014). One of these factors is likely the annual variability in larval survival. Based on our model, larval survival can vary 2.6-fold among years, which accounts for 30% of annual recruitment variability (8.3-fold) for ABT over the same time period as our simulation (ICCAT, 2017). This result highlights the need to consider yearly ecosystem states for the management of ABT.

Ocean models are well-suited for evaluating larval mortality for species like ABT because their dispersal is strongly influenced by large-scale ocean circulation which is well resolved by hydrodynamic models, their pelagic larval duration is short, and their low-trophic-level food is strongly influenced by bottom-up forcing. ABT is thus an optimal candidate for including ocean model-derived indices into stock assessments. The ocean modeling framework implemented in this study also allows for real-time and future predictions of larval survival, which could be used to inform assessment models on recruitment. Such predictions could be made with forecasts by hydrodynamic ocean models, like those already in operation by entities such as the National Oceanic and Atmospheric Administration. Furthermore, fishery-independent age-0 indices could be derived from the survival predicted from models like BLOOFINZ-IBM or simpler food limitation and predation maps like those presented in this study.

In future studies, estimates of survival could be compared to stock-recruitment data and abundances from SEAMAP surveys as a way to further refine the model. Additional realism could be included by initializing particles based on survey data or by incorporating the impact of maternal effects such as initializing egg weights based on the condition of spawning females. Because ABT are selective feeders, even within mesozooplankton size class (Shiroza *et al.,* this issue), added realism may also be achieved by combining NEMURO-GoM with a zooplankton food web model (Stukel *et al.,* this issue).

## CONCLUSIONS

Multiple hypotheses have been formulated to explain why ABT spawn in the GoM. Other regions in the Atlantic and Caribbean Seas contain similar conditions to the GoM (e.g. warm oligotrophic water), yet show no evidence of large-scale spawning. Our model shows that predation and starvation are equally important as mortality sources for larval ABT, though their magnitude and relative importance varies spatially and with larval age. Our results unequivocally support the hypothesis that the primary advantage of spawning in oligotrophic regions is to reduce predation risk. They further indicate that the GoM is an ideal spawning ground because of the region’s large shelf and strong mesoscale activity which together increases the chance of shelf water entrainment into offshore regions that may be crucial for ensuring both low starvation during the critical period and low predation later in development.

## ACKNOWLEDGEMENTS

Authors acknowledge collaboration with the Spanish Institute of Oceanography and ECOLATUN (CTM-2015-68473-R MINECO/FEDER) project.

## FUNDING

This paper is a result of research supported by the National Oceanic and Atmospheric Administration’s RESTORE Science Program under federal funding opportunity NOAA-NOS-NCCOS-2017-2004875, a NASA IDS grant #80NSSC17K0560, NSF Biological Oceanography grant #1851347, the NOAA Office of Education - Educational Partnership Program award NA16SEC48100009 (NOAA Center for Coastal and Marine Ecosystems), the NOAA NMFS Fisheries and the Environment program, and the Northern Gulf Institute (projects 18-NGI3-41 and 18-NGI3-52) under NOAA award NA16OAR4320199, and in part by the Cooperative Institute for Marine and Atmospheric Studies (CIMAS), a Cooperative Institute of the University of Miami and NOAA NA20OAR4320472.

## DATA ARCHIVING

Data from field-collected larvae presented here have been submitted to the National Oceanic and Atmospheric Administration’s (NOAA) National Centers for Environmental Information (NCEI) data repository and will also be archived at BCO-DMO (Biological and Chemical Oceanography Data Management Office) site https://www.bco-dmo.org/program/819631.

## FIGURE LEGENDS

**Fig 1**. (A) Relationship between larval length (mm) and prey length (mm) from gut content analysis of 255 individuals collected in the GoM (Shiroza *et al.,* this issue). Upper and lower bounds of prey size are shown in red. Blue dotted line defines the break between zooplankton (SZ, 0.02-0.2 mm) and large zooplankton (LZ, 0.2-1 mm) NEMURO-GoM state variables. (B) Relationship between larval weight (mg DW) and age (days post hatch) for individuals collected in the GoM. (C) Starvation as a function of an individual’s condition where ≥1.0 indicates ideal condition. Condition below 0.25 is used as a threshold for the “point-of-no-return” where larvae experience irreversible starvation (increased to 1.0 d^−1^ (not shown)). (D) Predation on egg and larvae as a function of simulated large mesozooplankton (e.g. PZ, 1-5 mm) biomass and example curves of individual length at 1, 2, 4, and 8 mm.

**Fig 2**. (A) Comparisons of mesozooplankton dietary contribution (% of total diet) as a function of larval length (mm) between field-collected (black) and simulated larvae (red). Whiskers extend to the 95% confidence interval. Outliers are denoted by (+) for observations and outliers for model are not shown. (B) Comparison of larval weight (mg DW) as a function of age (days post hatch) between field-collected larvae (black dots) and simulated larvae. Red line denotes model average with the 95% CI represented by shaded area. Dashed blue line denotes the average age simulated larvae reach postflexion.

**Fig 3**. (A) Survival as a function of time (days post spawn) with red solid lines denoting the range when individuals finish the egg stage, blue lines denoting the range when individuals finish yolk sac stage, and grey shaded region highlights the critical period predicted by the model. (B) Survival of larval tuna estimated for each year as a function of age (days post hatch) for each year (1993-2012) and black is mean of all years.

**Fig 4**. (A) Mortality a function of age with total (red), starvation (black), and predation (blue) plotted by separate lines. (B) Spatial variability of average starvation (d^−1^) and (C) predation (d^−1^) prior to early postflexion (i.e. 10 dph) computed by organizing particles within 0.12° × 0.12° spatial bins.

**Fig 5**. Spatial variability in larval survival to early postflexion (A,C) and late postflexion (B,D). Survival is computed by organizing particles based on their positons within 0.12° × 0.12° bins (A,B) and based on their spawning location (C,D).

**Fig 6.** (A) Survival (%) as a function of age for larvae located inside anticyclonic regions (ACR), cyclonic regions (CR), anticyclonic boundary regions (ACBR), cyclonic boundary region (CBR), common water (CW), and shelf region (SR). (B) Starvation (d^−1^) (C) and predation (d^−1^) as functions of age for the same regions as above.

**Fig. 7**. Mean and instantaneous food limitation index maps (A-D) and predation maps (E-J) for the month of May. Average food limitation index map for (A) first-feeding larvae (i.e. 3 dph) and (B) early postflexion (i.e. 10 dph). Instantaneous food limitation index map on May 15^th^ 1996 for (C) first-feeding larvae and (D) early postflexion. Average predation for (E) first-feeding larvae, (F) early postflexion larvae, and (G) late postflexion larvae (i.e. 17 dph). Instantaneous predation map on May 15^th^ 1996 for (H) first-feeding larvae, (I) early postflexion larvae, and (J) late postflexion larvae.

## Online Appendix 1 Derivation of larval fish sensory radius and predation-induced mortality

### Larval fish sensory radius

Ingestion is highly sensitive to sensory radius. To estimate sensory radius of larval tuna in our model we utilize a recently determined anatomical relationship for the visual acuity of larval tuna (Hilder *et al.*, 2019) along with a theoretical model of visual predation derived by Aksnes and Utne (1997) to account for the impact of light and water clarity.

First, we obtain the maximum reactive distance (MRD) for larval tuna from the minimum separable angle (MSA) using the relationship MSA = 4.699*L_lt_^−1.129^ (Hilder *et al.*, 2019), where L_lt_ is larval length. The MSA takes into account the density of light sensors on the retina (e.g. cones) along with the geometry of the eye lens to determine the minimum angle which can be distinguished by a predator. The MRD for a prey item of a given size (L_p_) can then be determined using (eqA1).

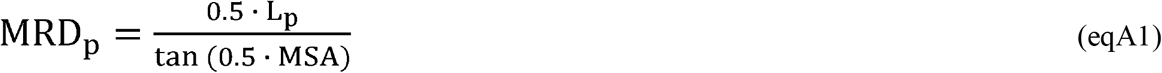

The reactive distance using this method serves as a theoretical anatomical maximum. To include the impact of light availability and water clarity on the reactive distance of larval tuna we utilize a theoretical model of aquatic visual predation developed by Aksnes and Utne (1997) (eqA2). The model assumes that a predator (in this case larval tuna) can detect a prey only if the difference between the retinal irradiance flux produced by a prey image and background is greater than some threshold. Unlike the minimum separable angle approach, the Aksnes and Utne (1997) model includes the impact of prey size as well as contrast to calculate reactive distance.

The Aksnes and Utne (1997) model includes a saturating light term where I_m_ is the maximum possible irradiance the visual system of a predator can process, I_z_ is the ambient light for a given depth, and k_i_ is the light half saturation coefficient, which was determined experimentally by Aksnes and Giske (1993). The light limitation term is multiplied by the prey contrast (C_p_), attenuation of the contrast signal across the reactive distance (D_p_ (in meters)) with beam attenuation (c). A_p_ is the area of the prey image and ΔE is the sensitivity threshold of a predator for detecting differences in retinal irradiance flux between a prey image and background. For the full derivation of (eqA2) see Aksnes and Giske, (1993) and Aksnes and Utne, (1997).

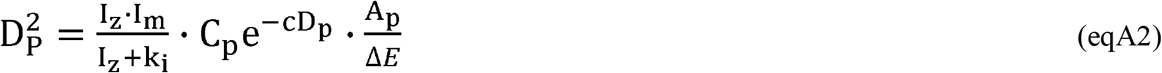

Since we are only interested in quantifying the impact of light, all terms which are not influenced by light availability or water clarity can be combined into a single constant X_1_. The constant can be solved for by assuming light saturated conditions (I_k_>>k_i_) with a prescribed beam attenuation and reactive distance in clear water. Beam attenuation is related to light attenuation (k_tot_) using the equation c=4 *k_tot_ (Aksnes and Giske, 1993) and in clear water k_tot_ is assumed to equal 0.04. The reactive distance is set using the MRD calculated from (eqA1). Once X_1_ has been determined the reactive distance with varying light and water clarity is obtained by iteratively solving (eqA3) using D_p_= MRD as an initial condition.

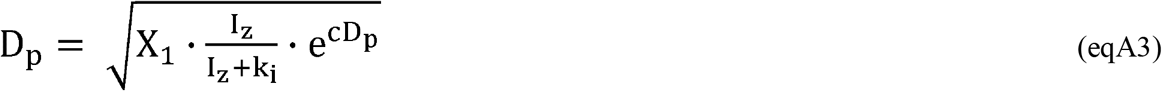

Ambient light levels (I_z_) are obtained from NEMURO-GoM which is forced by daily averaged surface shortwave irradiance fields estimated by the CFSR (Climate Forecast System Reanalysis) atmospheric model (see Shropshire *et al.,* 2020 for more details). The impact of water clarity is obtained using the total light attenuation term estimated from NEMURO-GoM. Vertical light attenuation in NEMURO-GoM is modeled using Beer’s law where k_tot_ is a function of attenuation due to water as well as small and large phytoplankton biomass.

Assuming a conical field of view with a given angle one can obtain the sensory radius from the estimated reactive distance in (eqA3) with the equation S_p_ = D_p_ · tan (Ø · 0.5). The location of eyes in pelagic larval fish are often located towards the top of the head with little to no ability to see behind them resulting in a maximum visual field of view angle of approximately 180°.

However, given the upward orientation of the eyes in larval tuna here we assume individuals perceive half of the possible perception field (e.g. Ø = 90°) which results in the reaction distance and sensory radius being equal (S_p_ = D_p_). As stated previously the MRD estimated in (eq3) is considered a theoretical maximum. Laboratory feeding studies have shown evidence suggesting the maximum behavioral reactive distance is approximately half of the anatomical MRD (Parker *et al.*, 2017). Hence, the reactive distance estimated in (eqA3) is first divided by a factor of 2.0 before calculating the sensory radius used to calculate ingestion with (eq2).

### Predation module derivation

Unlike starvation, individuals experience predation at all stages of the life cycle. Here we assume that predation on larval ABT is only a function of larval size. That is, the escape response is assumed to increase in proportion to the predator’s capture ability. We utilize the mortality rate on large mesozooplankton (PZ, 1 – 5 mm) estimated by NEMURO-GoM to approximate predation on larval tuna given the overlap in size. In NEMURO-GoM, mortality on PZ is modeled with a quadratic formulation which is often used in biogeochemical models to represent implicit loss terms on the highest trophic level due to an un-modeled predator that covaries in abundance with its prey. Specific mortality of PZ (M_PZ_) in NEMURO-GoM is expressed using (eq8), where m is the mortality rate parameter (d^−1^) as a function of temperature.

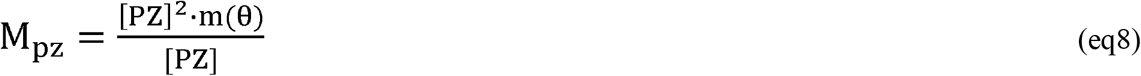

However, we could also write a mechanistic equation relating PZ mortality to the abundance of PZ, the abundance of an unmodeled predator (F) with a sensory radius (S_pz_) and swimming speed (v) (e.g., similar to (eq2), but for predator “F” feeding on PZ rather than larval ABT feeding on SZ and LZ). The concentration of PZ cancels and the equation can be written in terms of predator abundance and encounter rate (eq9).

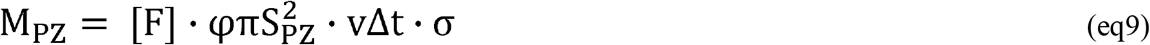

The same equation to (eq9) can be written for the specific mortality for larval tuna (M_LT_) where the predator now has a sensory radius (S_LT_). If we assume that the predators of PZ and larval ABT are broadly similar (a reasonable assumption, because of their overlap in size) the ratio of predation on larval tuna to PZ mortality is found to equal the ratio of their sensory radius squared. Predation on larval tuna can then be estimated using (eq10) where M_PZ_ is spatiotemporally varying as estimated by NEMURO-GoM.

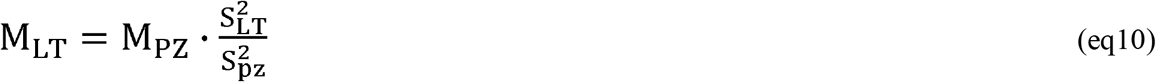

To determine S_LT_ and S_PZ_ we again start with the Aksnes and Utne (1997) visual predation model used to determine sensory radius for larval tuna. First, the reactive distance for a predator feeding on PZ is written as in (eq4). All terms that are not associated with reactive distance or prey attributes can be aggregated into a single constant X_2_ (eq11). The underlying assumption is that the influence of light availability and water clarity will have the same impact on the ability of a predator to capture PZ or larval ABT.

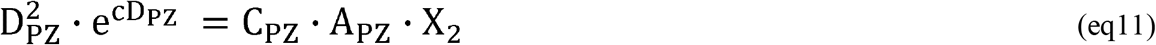

Similarly, (eq11) can be written for the reactive distance of a predator feeding on larval tuna (D_LT_) using the prey contrast (C_LT_) and area (A_LT_). Taking the two equations, D_PZ_ can be written in terms of larval tuna attributes using two parameters (β, γ) that relate PZ size and contrast to larval tuna size and contrast.

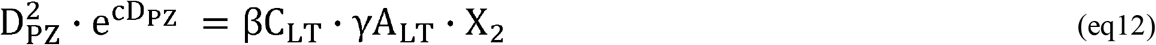

The left hand side of (eq11) (with subscripts LT) can be substituted on the right hand side of (eq12) resulting in a single equation that relates the reactive distance of a predator feeding on PZ and larval tuna (eq13).

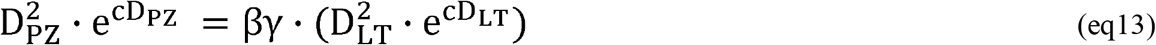

Like most mesozooplankton, larval tuna are highly transparent. Here we assume that the contrast of PZ and larval ABT is equal (β = 1.0) and hence the term can be removed. The prey area scaling term can be expanded (i.e. γ = (π·0.5·L_PZ_)^2^/(π·0.5·L_LT_)^2^) and solving for the reaction distance of a predator feeding on larval tuna results in (eq14).

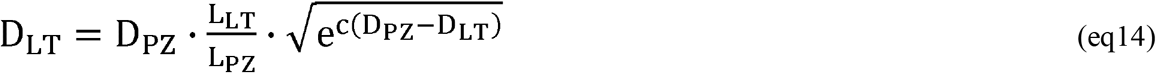

The equation can be solved iteratively by prescribing a value for D_PZ_ and assuming for an initial condition that the reactive distance of a predator feeding on PZ and larval ABT is equal (D_LT_ = D_PZ_). Here D_PZ_ is approximated using three assumptions: (1) PZ is on average 2.0 mm in length (PZ in NEMURO-GoM is defined as 1-5 mm); (2) the predator feeding on PZ has an average predator to prey ratio (PPR) = 10; and (3) the predator has a reactive distance (D_P_) of 2 body lengths, resulting in D_PZ_ = 40 mm. Assumptions (2) and (3) do not influence the final predation rate on larval tuna because after expanding D_PZ_ in (eq14) (i.e. D_PZ_ = L_PZ_ · PPR · D_P_) one finds that L_LT_ is also scaled by both PPR and D_p_ terms. Since the sensory radius is equal to reactive distance when assuming a 90° field of view the PPR and D_p_ terms cancel when computing the ratio S_LT_/S_PZ_. Finally, the sensory radius for PZ and LT are inserted into (eq8) to calculate spatiotemporally-varying predation on LT which gives (eq15).

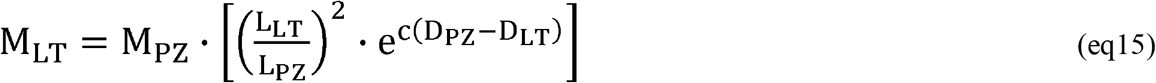

We note that the exponential term is found to be nearly constant (expression inside the exponential is on the order −1×10^−2^ to −1×10^−3^) resulting in predation on LT increasing approximately in proportion to the ratio of larval and PZ length squared.

During model tuning the predation rate was adjusted using values of 2 – 4 mm for the average size of PZ. We found that L_PZ_ of 2 mm resulted in realistic values of mortality when added with starvation (Houde, 2002). For predation during the egg stage L_LT_ is set to 1.0 mm while L_LT_ for yolk-sac and post yolk-sac larvae is set based on the previously described length to age relationship. During the egg and yolk-sac stage larvae experience lower rates of predation than mortality on PZ estimated by NEMURO-GoM given their smaller size. Mortality on larval ABT is equal to the mortality on PZ when larval ABT are the same size as PZ (i.e. 2 mm) and increases non-linearly after this point.

## Notes

### Competing Interest Statement

The authors have declared no competing interest.

