## Supplemental for "Trade-offs between risks of predation and starvation in larvae make the shelf break an optimal spawning location for Atlantic Bluefin tuna"

**S1	BLOOFINZ-IBM Parameters**

**Table S1:** Model parameters description and values referenced in equations and parameter sensitivity experiments.

| Parameter | | | Units | Value |
| --- | --- | --- | --- | --- |
| **Tunable Model Parameters** | | | | |
| **Symbol** | **Name** | **Description** |  |  |
| *V* | SwmSpd | Cruise swimming speed | Body length s^-1^ | 2.0 |
| *Φ* | Vis2DFac | Two dimensional field of view fraction | Non-Dim. | 0.5 |
| *ϵ* | GrossGrowthEff | Gross growth efficiency | % Ingestion | 0.3 |
| *Α* | AssimEff | Absorption efficiency | % Ingestion | 0.7 |
| *σ_p_* | CaptSucc | Capture success | Non-Dim. | 0.75 |
| *t_c_* | Q10Temp | Metabolic requirement temperature dep. | °C^-1^ | 0.0693 |
| *t_c_* | Q10Temp | Yolk-sac temperature dep. | °C^-1^ | 0.0693 |
| *t_c_* | Q10Temp | Growth (in length) temperature dep. | °C^-1^ | 0.0693 |
| *t_c_* | Q10Temp | Gut turnover temperature dep. | °C^-1^ | 0.0693 |
| *L_PZ_* | PZlen | Average length of Predatory Zoo. (PZ) | mm | 2.0 |
| *c_f_* | CarbFac | Carbon fraction | Non-Dim. | 0.4 |
| ‘’ | StarvMax | Maximum starvation rate | d^-1^ | 0.3 |
| ‘’ | PONRmort | Point-of-no return starvation rate | d^-1^ | 1.0 |
| ‘’ | PONRfrac | Point-of-no-return threshold | % Body mass | 0.25 |
| ‘’ | GutFull | Gut fullness | % Body mass | 0.1 |
| ‘’ | GutTurn | Gut turnover time at 26°C | hours | 3.0 |
| ‘’ | BasePredMort | Minimum predation rate | d^-1^ | 0.05 |
| ‘’ | ExogFeedAge | Age at exogenous feeding at 26°C | Days post hatch | 2.0 |
| ‘’ | LarvLightHalfSat | Aksnes et al visual light parameter | Watts m^-2^ | 1.0869 |
| ‘’ | MRD2BRD | Max Reactive Dist / Behav React Dist | Non-Dim. | 2.0 |
| ‘’ | EggDiam | Egg diameter | mm | 1.0 |
| ‘’ | EggDiamSTD | Egg diameter standard deviation | mm | 0.04 |
| ‘’ | FieldSampleTmpAvg | Assumed water temp of collected larvae | °C | 26 |
| ‘’ | TmpAdjHatchProb | Temperature adjustment Equation 2 | °C | -2 |
| ‘’ | PostWeightThresh | Postflexion weight | mg DW | 0.54 |
| **Fitted Relationship Parameters** | | | | |
| **Symbol** | **Name** | **Description** |  |  |
| ‘’ | HatchTime_p1 | Equation S1 – Parameter 1 |  | 4.66 |
| ‘’ | HatchTime_p2 | Equation S1 – Parameter 2 |  | -0.11 |
| ‘’ | HatchProb_p1 | Equation S2 – Parameter 1 |  | -1.27 |
| ‘’ | HatchProb_p2 | Equation S2 – Parameter 2 |  | 63.78 |
| ‘’ | HatchProb_p2 | Equation S2 – Parameter 3 |  | 727.98 |
| ‘’ | Age2Weight_p1 | Equation S3 – Parameter 1 |  | 0.0547 |
| ‘’ | Age2Weight_p2 | Equation S3 – Parameter 2 |  | 0.2157 |
| ‘’ | Age2Weight_std_p1 | Standard Deviation (Age2Weight_p1) |  | 0.0052 |
| ‘’ | Age2Weight_std_p2 | Standard Deviation (Age2Weight_p2) |  | 0.0108 |
| ‘’ | max_Age2Weight_p1 | Equation S4 – Parameter 1 |  | 0.1128 |
| ‘’ | max_Age2Weight_p1 | Equation S4 – Parameter 2 |  | 0.1714 |
| ‘’ | Age2Length_p1 | Equation S5 – Parameter 1 |  | 0.4433 |
| ‘’ | Age2Lenght_p2 | Equation S5 – Parameter 2 |  | 1.8590 |
| ‘’ | Age2Lenght_std_p1 | Standard Deviation (Age2Weight_p1) |  | 0.011 |
| ‘’ | Age2Length_std_p2 | Standard Deviation (Age2Weight_p2) |  | 0.0802 |
| ‘’ | upp_LarvLen2PreyLen_p1 | Equation S6 – Parameter 1 |  | 0.071 |
| ‘’ | upp_LarvLen2PreyLen_p2 | Equation S6 – Parameter 2 |  | 0.22 |
| ‘’ | low_LarvLen2PreyLen_p1 | Equation S7 – Parameter 1 |  | 0.015 |
| ‘’ | low_LarvLen2PreyLen_p2 | Equation S7 – Parameter 2 |  | 0.04 |
| ‘’ | MinSepAng_p1 | Equation S8 – Parameter 1 |  | 4.699 |
| ‘’ | MinSepAng_p2 | Equation S8 – Parameter 2 |  | -1.129 |

**S2	Parameter sensitivity**

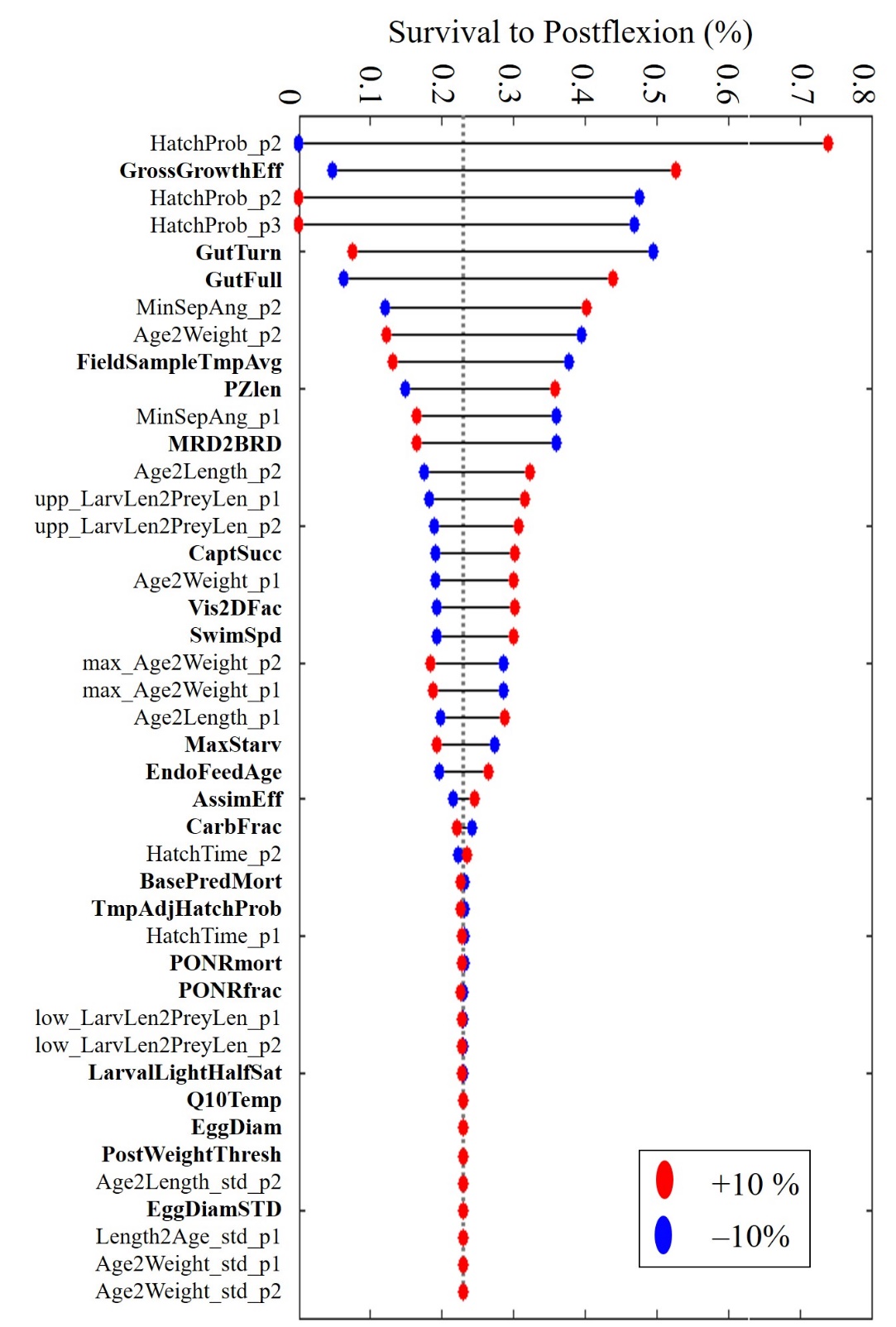

**Figure S1:** Survival to postflexion for 43 independent parameter sensitivity experiments which included -10% (blue) /+10% (red). Bolded parameters denote the main tunable model parameters while non-bolded text indicates parameters that are associated with fitted relationships derived from field data.

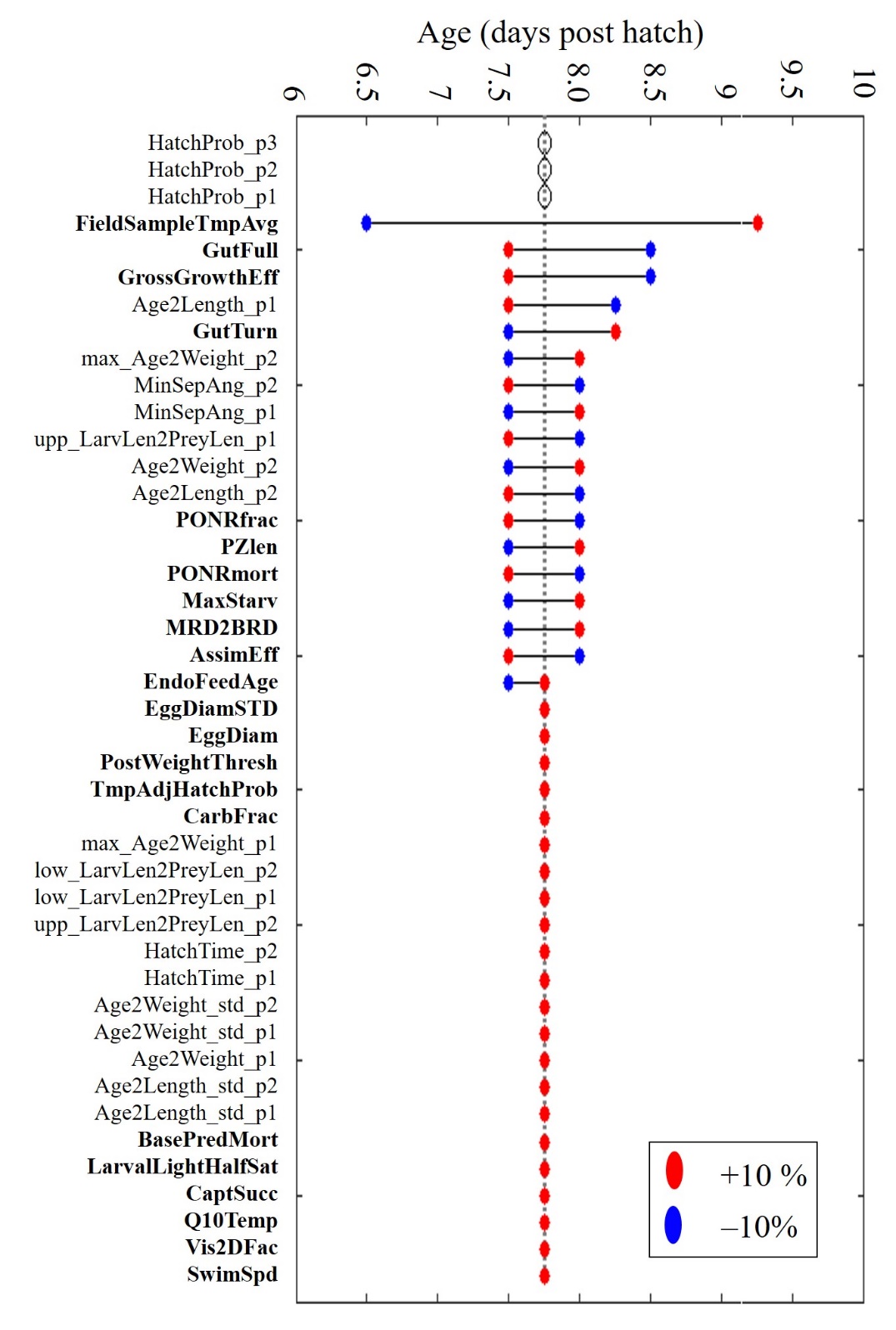

**Figure S2:** Age at which starvation and predation become approximately equal calculated for 43 independent parameter sensitivity experiments which included -10% (blue) /+10% (red). Bolded parameters denote the main tunable model parameters while non-bolded text indicates parameters that are associated with fitted relationships derived from field data.

**S3	Model diagnostics as a function of larval age**

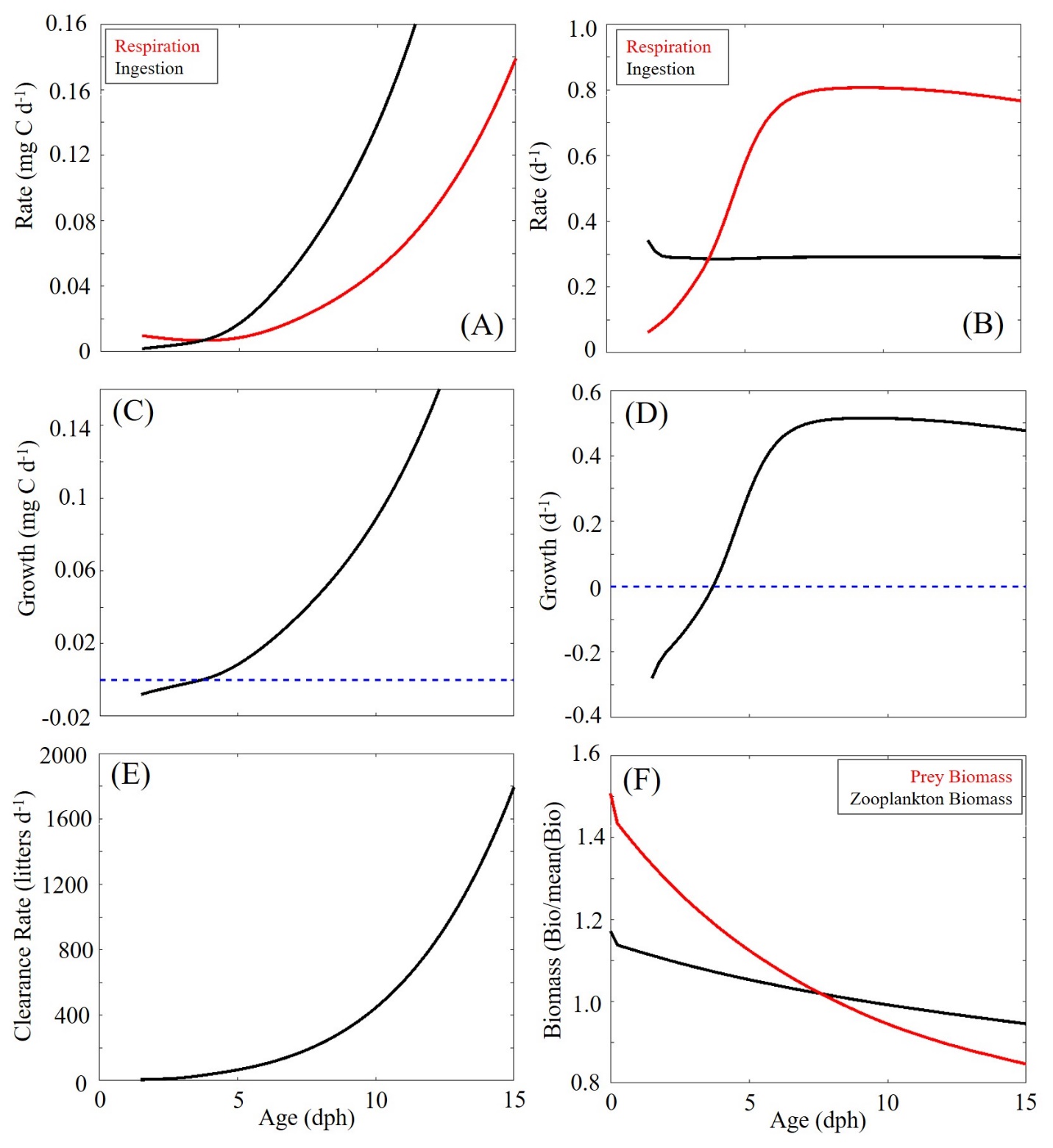

**Figure S3:** (A) Larval respiration and ingestion (mg C d^-1^) as a function on larval age (days post hatch). (B) Respiration and ingestion (d^-1^) normalized to larval weight as a function of larval age. (C) Larval growth and (D) specific growth as a function of larval age. (E) Clearance (liters d^-1^) as a function of larval age. (F) Zooplankton and prey biomass normalized to their respective average biomass.

**S4	Fitted relationships used in BLOOFINZ-IBM**

*Egg stage (required hours to hatch):*

$h=4.66 \cdot exp(-0.11\cdot\theta)$	(eq. S1)

*Hatch probability*

$x=-1.27\cdot\theta^{2}+63.78\cdot\theta-727.98$	(eq. S2)

*Age to weight relationship*

$W=0.0547 \pm0.0052 \cdot exp(0.2157 \pm0.0108 \cdot A)$	(eq. S3)

*Maximum age to weight relationship*

$W=0.1128 \cdot exp(0.1714 \cdot A)$	(eq. S4)

*Age to length relationship*

$L=(0.4433 \pm0.0111\cdot A+1.8590\pm0.0802)$	(eq. S5)

*Larval length to prey length (upper bound)*

$L_{P}=0.071\cdot L_{T}+0.22$	(eq. S6)

*Larval length to prey length (lower bound)*

$L_{P}=0.015\cdot L_{T}+0.04$	(eq. S7)

*Minimum Separable Angle*

$MSA= 4.699 \cdot L_{\mathrm{LT}}^{-1.129}$	(eq. S8)
